# The DNA-binding protein HTa from *Thermoplasma acidophilum* is an archaeal histone analog

**DOI:** 10.1101/564930

**Authors:** Antoine Hocher, Maria Rojec, Jacob B. Swadling, Alexander Esin, Tobias Warnecke

## Abstract

Histones are a principal constituent of chromatin in eukaryotes and fundamental to our understanding of eukaryotic gene regulation. In archaea, histones are phylogenetically widespread, often highly abundant, but not universal: several archaeal lineages have lost histone genes from their coding repertoire. What prompted or facilitated these losses and how archaea without histones organize their chromatin remains largely unknown. Here, we use micrococcal nuclease digestion followed by high-throughput sequencing (MNase-Seq) to elucidate primary chromatin architecture in an archaeon without histones, the acido-thermophilic archaeon *Thermoplasma acidophilum*. We confirm and extend prior results showing that *T. acidophilum* harbours a HU family protein, HTa, that is highly expressed and protects a sizeable fraction of the genome from MNase digestion. Charting HTa-based chromatin architecture across the growth cycle and comparing it to that of three histone-encoding archaea (*Methanothermus fervidus, Thermococcus kodakarensis* and *Haloferax volcanii*), we then present evidence that HTa is an archaeal histone analog. HTa-protected fragments are GC-rich, display histone-like mono- and dinucleotide patterns around a conspicuous dyad, exhibit relatively invariant positioning throughout the growth cycle, and show archaeal histone-like oligomerization dynamics. Our results suggest that HTa, a DNA-binding protein of bacterial origin, has converged onto an architectural role filled by histones in other archaea.

## Introduction

Across all domains of life, DNA is intimately associated with proteins that wrap, package, and protect it. Bacteria typically encode multiple small basic proteins that are dynamically expressed and fulfil a variety of architectural roles by bridging, wrapping, or bending DNA (Dillon and Dorman 2010). Some of these proteins are phylogenetically widespread, others restricted to specific lineages (Swiercz *et al*. 2013; Lagomarsino *et al*. 2015). Bacterial chromatin, on the whole, is diverse and dynamic over both physiological and evolutionary timescales. In contrast, a single group of proteins has come to dominate chromatin in eukaryotes: histones. Assembling into octameric complexes that wrap ~147bp of DNA, eukaryotic histones not only mediate genome compaction but also establish a basal landscape of differential accessibility, elaborated via a plethora of post-translational modifications, that is fundamental to our understanding of eukaryotic gene regulation (Jiang and Pugh 2009; Bai and Morozov 2010).

Histones are also widespread in archaea (Adam *et al*. 2017; Henneman *et al*. 2018). They have the same core fold (Decanniere *et al*. 2000; Mattiroli *et al*. 2017) as eukaryotic histones, but lack N- and typically also C-terminal tails, the principal substrates for post-translational modifications in eukaryotes (Henneman *et al*. 2018). Dimers in solution, they assemble into tetramers that wrap ~60bp of DNA (Bailey *et al*. 1999). This minimal nucleosomal unit can be extended, at least in some archaea, into a longer oligomer via incorporation of additional dimers (Xie and Reeve 2004; Maruyama *et al*. 2013; Mattiroli *et al*. 2017). Like their eukaryotic counterparts, archaeal nucleosomes preferentially bind sequences that, for example by means of periodically spaced GC/GG/AA/TT dinucleotides, facilitate wrapping (Bailey and Reeve 1999; Bailey *et al*. 2000). On average, nucleosome occupancy is higher on more GC-rich sequences and lower around transcriptional start and end sites (Ammar *et al*. 2011; Nalabothula *et al*. 2013), which tend to be relatively AT-rich. The precise role of archaeal histones in transcription regulation, however, remains poorly understood (Gehring *et al*. 2016).

Although widespread, archaeal histones are less phylogenetically entrenched than histones in eukaryotes: they have been deleted experimentally in several species without dramatic effects on transcription and growth (Heinicke *et al*. 2004; Weidenbach *et al*. 2008; Dulmage *et al*. 2015) and lost altogether from a handful of archaeal lineages (Adam *et al*. 2017). A particularly intriguing case concerns the thermophilic acidophile *Thermoplasma acidophilum*, which lacks histone genes but instead encodes a protein named HTa (**H**istone-like protein of ***T**hermoplasma acidophilum*). Based on its primary sequence (DeLange *et al*. 1981; Drlica and Rouvière-Yaniv 1987) and predicted secondary, tertiary and quaternary structure (Figure 1a-b), HTa is a member of the HU family of proteins, which are broadly distributed across bacterial phyla (Table S1) and often abundant constituents of bacterial chromatin. This includes *Escherichia coli* (Figure 1c), where the average cell during exponential growth contains an estimated 30,000-55,000 HU molecules (Azam *et al*. 1999). Although individual members of the HU family have diverged in their DNA binding properties, even distant homologs display functional similarities, constraining negative supercoils and binding not only B-form but also structurally unusual DNA such as cruciforms (Grove 2011). Outside bacteria, HU proteins are relatively rare. They are found with a modicum of phylogenetic persistence only in some single-celled eukaryotes (Figure 2a , Supplementary Text), where they are known to play important functional roles (Sasaki *et al*. 2009; Gornik *et al*. 2012), and in a single clade of archaea: the Thermoplasmatales/deep-sea hydrothermal vent euryarchaeota (DHVE2 group). Phylogenetic reconstruction of the HTa/HU gene family suggests that HTa was acquired via horizontal gene transfer from bacteria at the root of this clade. However, HTa has sufficiently diverged from its bacterial homologs – and the event is sufficiently ancient – to prevent facile identification of a specific bacterial donor clade (Figure 2, Supplementary Text).

**Figure 1.**
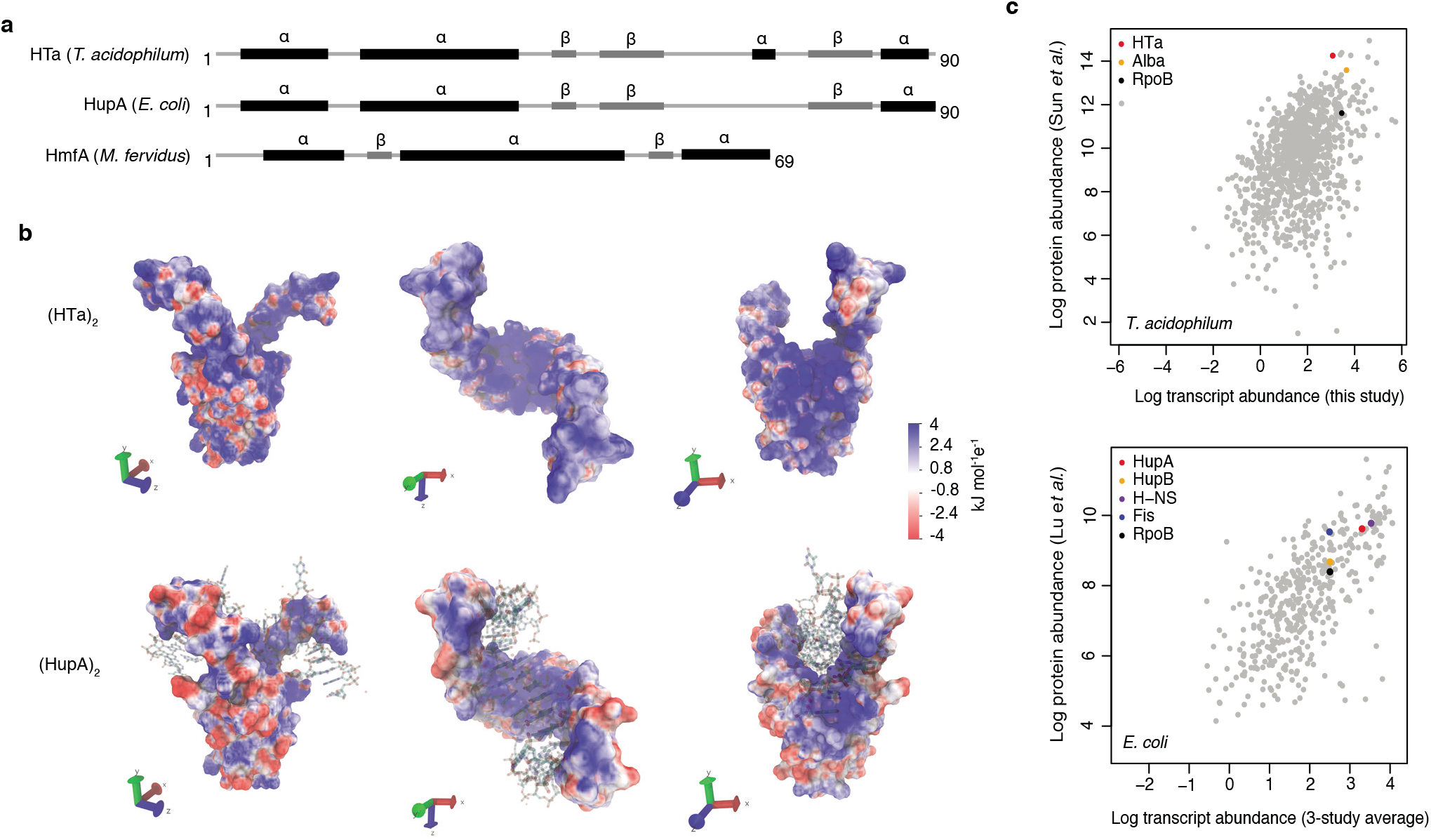
Predicted structure and measured abundance of HTa. **(a)** Predicted secondary structures of HTa (*T. acidophilum*), the bacterial HU protein HupA (*E. coli*), and the archaeal histone protein HmfA (*M. fervidus*). **(b)** Predicted quaternary structure of the (HTa)_2_ homodimer compared to the crystal structure of (HupA)_2_ (PDB: 1p51) bound to DNA. Colour gradients represent charge densities mapped onto the solvent accessible surface area of (HTa)_2_ and (HupA)_2_. Note the extended patches of stronger positive charge for (HTa)_2_ compared to (HupA)_2_, particularly in the stalk region. **(c)** Correlation of transcript and protein abundances for *T. acidophilum* and *E. coli*. HTa and HU are highlighted along with some additional chromatin-associated proteins. Data sources: *T. acidophilum* protein abundance: Sun et al. (2010); *E. coli* protein abundance: Lu et al. (2007). *E. coli* transcript abundance is an average across three previous studies as reported by Lu et al. (2007).

**Figure 2.**
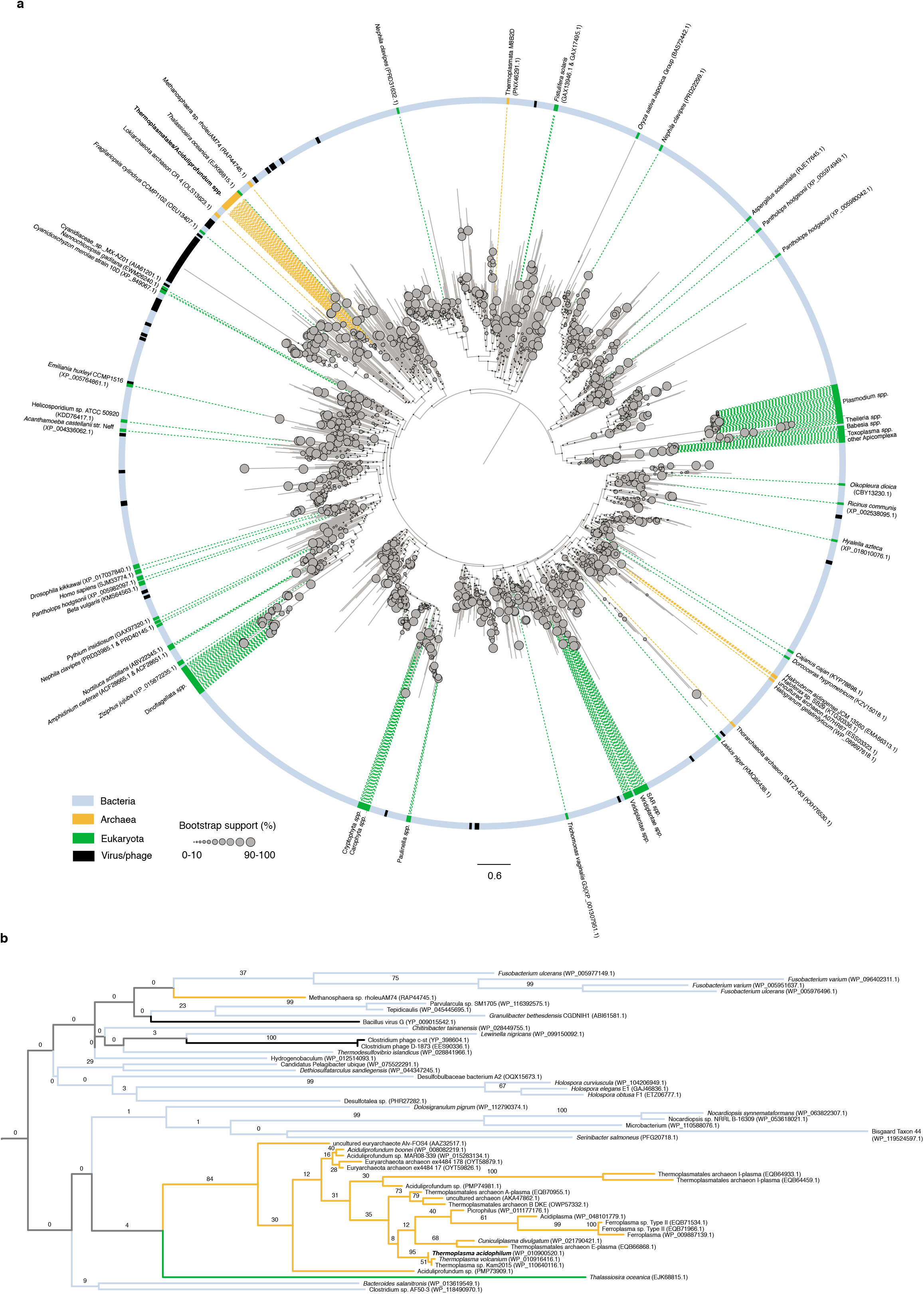
Phylogenetic relationships of HU family proteins from bacteria, eukaryotes, and archaea. **(a)** protein-level phylogenetic tree of HU proteins including HTa (see Methods for details on phylogenetic reconstruction). The tree is midpoint-rooted. Reported domain-level membership (Bacteria, Archaea, etc.) of different proteins is colour-coded in the outer circle and on the dotted lines that point to individual branches. See Supplementary Text for a critical evaluation of domain assignments and likely assembly contaminants. Bootstrap support values (%) for individual branches, visually encoded as node diameters, illustrate poorly resolved relationships at deeper nodes. **(b)** Excerpt of the phylogeny shown above, highlighting good support (84%) for a monophyletic origin of HU proteins in the Thermoplasmatales/DHVE2 clade and their uncertain affiliation to other HU family members.

In *T. acidophilum*, HTa is highly abundant (Figure 1c) and protects ~25-35% of the genome from micrococcal nuclease (MNase) digestion (Searcy and Stein 1980; Thomm *et al*. 1982), consistent with a global role in structuring *T. acidophilum* chromatin. Pioneering work by Searcy and colleagues showed that MNase digestion of native *T. acidophilum* chromatin yields two distinct fragment sizes of ~40bp and ~80bp (Searcy and Stein 1980). The same authors observed an indistinguishable banding pattern when they digested calf thymus DNA following *in vitro* reconstitution with purified HTa, suggesting that native HTa is sufficient for and likely the principal mediator of protection from MNase digestion in *T. acidophilum* (Searcy and Stein 1980). It remains unknown, however, where HTa binds to the *T. acidophilum* genome *in vivo*; whether HTa binds in a sequence-specific manner or promiscuously; whether it requires particular post-translational modifications to carry out its functions; whether binding is dynamic in response to environmental changes; and how binding relates to functional genomic landmarks. Crucially, we do not know how HTa-mediated chromatin organization in *T. acidophilum* compares to that in histone-encoding archaea: do HTa and histones fill similar functional and architectural niches? Or are their binding patterns and functional repercussions entirely distinct?

Here, to begin to address these questions, we characterize genome-wide chromatin organization mediated by HTa in *T. acidophilum*. Using MNase treatment coupled to high-throughput sequencing, we find footprints of protection throughout the genome. Confirming prior results (Searcy and Stein 1980), we observe a bimodal distribution of protected fragment sizes and subsequently infer small and large binding footprints. The more common smaller footprints are well predicted by simple sequence features and display a general preference for GC-rich sequences. Their sequence preferences, positioning around transcription start sites, and static behavior throughout the growth cycle are reminiscent of archaeal histones rather than well-characterized bacterial HU homologs. In addition, we present evidence that larger fragments are frequently derived from nucleation-extension events, similar to what has been observed for archaeal histones that have the capacity to oligomerize (Maruyama *et al*. 2013; Nalabothula *et al*. 2013; Mattiroli *et al*. 2017). Our results suggest that, in key aspects of its molecular behaviour, HTa can be regarded as an archaeal histone analog.

## Results

### *Primary chromatin structure across the* T. acidophilum *growth cycle*

To elucidate genome-wide HTa binding *in vivo*, we carried out a series of MNase experiments across the *T. acidophilum* growth cycle. Throughout, MNase digestion yielded protected fragments of two distinct sizes (Figure 3a), in line with previous results (Searcy and Stein 1980). MNase digestion of *E. coli* cells expressing recombinant HTa produced a very similar protection pattern (Figure 3b). This demonstrates that HTa readily binds DNA outside its native cellular environment and does not require *T. acidophilum*-specific post-translational modifications or binding partners to protect from MNase digestion, consistent with previous *in vitro* reconstitution experiments (Searcy and Stein 1980). In contrast, native *E. coli* HU (HupA), even when strongly over-expressed from the same plasmid, does not confer significant protection under equivalent digestion conditions (Figure 3b). Neither do HU orthologs from *Thermobacillus composti* and *Lactobacillus floricola* (Figure S1a), which have higher sequence identity to HTa (37% and 39% compared to 27% for *E. coli* HupA). HTa is therefore unusual, although perhaps not unique (Ghosh and Grove 2004; Mukherjee *et al*. 2008) among HU family proteins in its capacity to protect DNA from MNase attack. Interestingly, the MNase profile of HTa-expressing *E. coli* (see Figure S1a in particular) suggests protection of additional, longer fragments, not as readily apparent in the *T. acidophilum* digest (but see Figure 3c), an observation to which we will return below.

**Figure 3.**
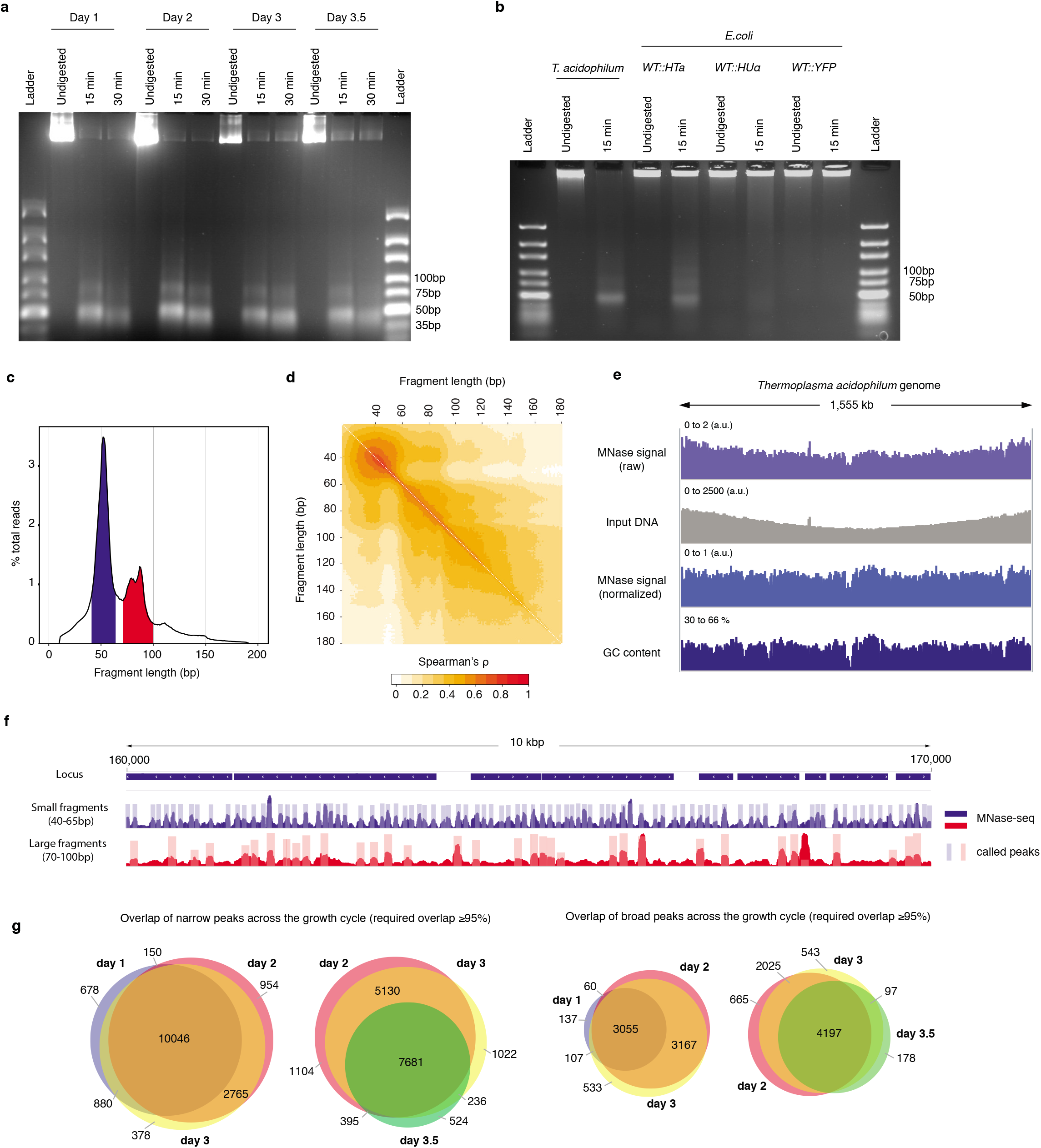
HTa-mediated primary chromatin architecture in *T. acidophilum* mapped by MNase-Seq. **(a)** Agarose gel of MNase digestion products from *T. acidophilum* sampled across the growth cycle. Growth phases are given as days after inoculation, digestion time in minutes. **(b)** Agarose gel of MNase digestion products from *T. acidophilum* (day 2) along with digestion products of *E. coli* ectopically expressing HTa, HupA or YFP (see Methods). **(c)** Distribution of the lengths of fragments mapped to the *T. acidophilum* genome (pooled across all four replicates from day 2), highlighting fragment size ranges that correspond to small (blue) and large (red) fragments, as defined in the main text. **(d)** Correlation matrix comparing genome-wide MNase-Seq coverage signal, computed at base pair resolution, between reads of defined sizes (pooled replicates, day 2). **(e)** Genome-wide MNase-Seq signal prior to and after normalization with sonicated DNA input (see Methods), along with GC content profile along the *T. acidophilum* chromosome, computed using a 51bp moving window. **(f)** Example of coverage and called peaks across a 10kb region of the *T. acidophilum* chromosome. **(g)** Overlap of detected narrow and broad peaks across the growth cycle. Note that different sections/overlaps are only qualitatively but not quantitatively proportional to absolute peak numbers.

Next, we sequenced the *T. acidophilum* DNA fragments that survived MNase treatment using Illumina paired-end technology (see Methods). As anticipated, two major fragment size classes are present across all stages of the growth cycle (Figure 3c, Figure S1b). For downstream analysis, we define small (large) fragments as 40-65bp (70-100bp) in size and note the following: first, the twin peaks centred around ~85bp (Figure 3c), separated by approximately one helical turn, were evident across biological replicates (Figure S1b). At present, we do not know whether this reflects discontinuous unwrapping and digest or the presence of distinct binding species. However, genome-wide occupancy of 70-80bp and 80-90bp fragments is highly correlated (Spearman’s ρ=0.76, P< 2e-16, Figure 3d) and we therefore consider 70-80bp and 80-90bp fragments jointly. Second, modal fragment sizes (~50bp and ~85bp) are slightly larger than those reported previously (~40bp and ~80bp) (Searcy and Stein 1980). At least in part, this reflects digestion conditions: fragments obtained after doubling digestion time from 15 to 30 minutes are shorter and map inside larger footprints (Figure S2). We chose and persisted with a somewhat milder digest here to avoid over-digestion of small fragments.

We then mapped fragments, irrespective of their size, to the *T. acidophilum* genome (see Methods). Protection across the genome is both ubiquitous and heterogeneous, with relatively even coverage along the origin-terminus axis once increased copy number of early replicating regions is taken into account (see Methods, Figure 3e-f). Major drop-offs in coverage correspond to areas of low GC content, as evident from Figure 3e, formally shown in Figure S3, and further explored below. For any given growth phase, genome-wide occupancy is highly correlated across replicates (mean ρ=0.89, P<2.2e-16 for all pairwise comparisons). For each time point, we therefore merged reads across replicates and called peaks independently for small and large fragments (see Methods). Globally, peak locations and occupancy vary little across the growth cycle (Figure 3g). Below, we will focus on data from exponential phase (day 2), where we observe 13,915 narrow and 6,887 broad peaks, before discussing variability of HTa-mediated chromatin architecture across the growth cycle in the context of transcriptional changes.

### Analysis of HTa binding footprints suggests histone-like oligomerization behaviour

Considering read coverage around called peaks, it is evident that small and large fragments often overlap (Figure 3f, Figure 4c,f), indicating the presence of different protective entities in the same location across cells. Importantly, broad peaks are typically a combination of small and large fragments and often show asymmetric coverage (caused by smaller fragments) around the summit of the inferred peak (see Figure 4a for an example). Given that some archaeal histones (Maruyama *et al*. 2013; Mattiroli *et al*. 2017) and bacterial HU proteins (Hammel *et al*. 2016; Hołówka *et al*. 2017) can form oligomers, we reasoned that asymmetric coverage might contain valuable information about the potential genesis of larger fragments from smaller nucleation sites. To retain this signal, lost when averaging over individual peaks in aggregate plots, we re-oriented the coverage signal as displayed in Figure 4b, revealing that smaller fragments are aligned to the edge – rather than the centre – of broad-peak footprints (Figure 4d,g).

**Figure 4.**
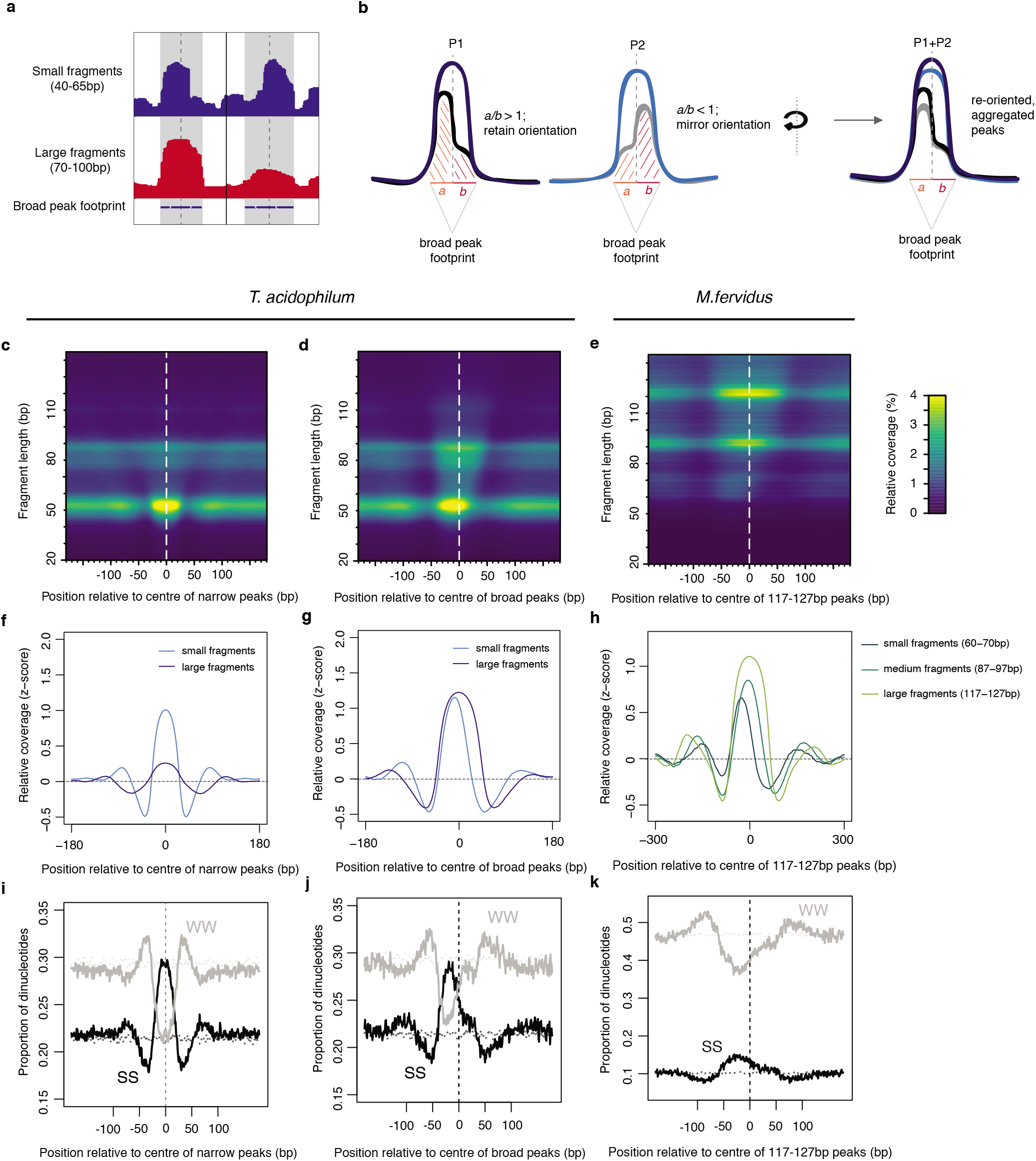
Asymmetric coverage signals around peaks in *T. acidophilum* and *M. fervidus* that track underlying nucleotide content. **(a)** Empirical example and **(b)** schematic describing our approach to re-orienting coverage signals at broad peaks based on the coverage of small fragments around the dyad axis. **(c, d)** Heat maps illustrating MNase-seq coverage by fragment length relative to the centre of narrow and broad peaks in *T. acidophilum*. Coverage around broad peaks is oriented as explained in (b). **(e)** Analogous heat map illustrating coverage by fragment length relative to the centre of large peaks (corresponding to the binding footprints of octameric histone oligomers) in *M.fervidus*. **(f, g, h)** Normalized coverage for *T. acidophilum* small (40-65bp) and large (70-100bp) fragments and *M. fervidus* fragment ranges corresponding to the expected footprint sizes of histone tetramers, hexamers, and octamers. **(i, j, k)** Proportion of SS (= CC|CG|GC|GG) and WW (= AA|AT|TA|TT) dinucleotides at the same relative positions as (c, d, e). Dotted lines indicate the proportion of SS or WW dinucleotides expected by chance, estimated via random sampling of 25000 regions of equal size in each genome.

We then applied the same procedure to MNase data we had independently generated for the histone-containing archaeon *Methanothermus fervidus* (see Methods), which – thanks to the work of Reeve, Sandman and others – is the source of much of our foundational knowledge about archaeal histones. Comparing *M. fervidus* to *T. acidophilum*, we find a very similar nested, edge-aligned structure of smaller fragments within broader peaks (Figure 4e,h). In *M. fervidus*, this pattern is consistent with oligomerization, in dimer steps, from the minimal histone tetramer (Maruyama *et al*. 2013; Mattiroli *et al*. 2017). Whether the pattern in *T. acidophilum* similarly reflects direct physical contact or is caused by closely adjacent binding of independent HTa complexes remains to be established. We note that our approach to retaining asymmetric coverage signals might have broader utility in characterizing oligomerization behaviour of DNA-binding proteins from MNase or similar high-resolution foot-printing data.

### HTa exhibits histone-like sequence preferences

Next, we asked what factors govern HTa binding in general and presumed nucleation-extension dynamics in particular. As the MNase signal broadly tracks GC content (Figure 3e), we first considered nucleotide enrichment patterns associated with peaks. At a coarse level, we find that both broad and narrow peaks exhibit relatively elevated GC content at their centre and are flanked by short stretches of GC depletion (Figure 4i,j). In line with promiscuous binding, a specific binding motif, as one would observe for most transcription factors, is not evident (Figure S4a). For broader peaks, it is worth noting that, in both *T. acidophilum* and *M. fervidus*, once we have re-oriented the small fragment coverage signal as described above, peak-internal GC content positively tracks small fragment abundance (Figure 4j,k), supporting a model where nucleation happens on more GC-rich sites whereas sequence need not be as favourable for subsequent extension events. Note, however, that while sequence might be more important for nucleation than extension, it is by no means irrelevant for the latter: for example, narrow peaks where large fragments are rare tend to be flanked by more AT-rich sequence in both *T. acidophilum* and *M. fervidus*, suggesting that sequence can prevent or at least predispose against oligomer formation (Figure S4b,c).

We then considered dinucleotide frequencies in reads of defined lengths, restricting the analysis to reads that overlap previously defined peaks by at least 90% and discarding duplicate reads that mapped to the same genomic location, in order to not bias results towards highly occupied footprints. Read-internal dinucleotide profiles confirm an overall GC preference but also reveal local enrichment/depletion patterns, notably a short central track of reduced GC enrichment (Figure 5a) in *T. acidophilum* as well as histone-encoding archaea, which is symmetric around the HTa/nucleosome dyad. Reassuringly, local enrichment patterns get weaker and eventually disappear when considering reads increasingly further away from modal fragment lengths (Figure S5). Unexpectedly, we also find mononucleotide and RR/YY (R=A or G; Y=T or C) profiles similar to those previously observed for – and attributed to the unique geometry of – eukaryotic histones (Reynolds *et al*. 2010; Ioshikhes *et al*. 2011; Quintales *et al*. 2015). These profiles show strong counter-phasing and asymmetry across the dyad (Figure 5b,c) and are particularly prominent in *T. acidophilum* and *T. kodakarensis*. These observations suggest that symmetric and asymmetric nucleotide enrichments across a dyad axis are not limited to nucleosomes and, furthermore, imply that the HTa-DNA complex is symmetric. All three observations – a general preference for GC-rich sequence, symmetric WW/SS and asymmetric mononucleotide/RR/YY frequencies around the dyad axis – are strongly reminiscent of archaeal as well as eukaryotic histones (Peckham *et al*. 2007; Kaplan *et al*. 2008; Tillo and Hughes 2009; Ammar *et al*. 2011; Nalabothula *et al*. 2013).

**Figure 5.**
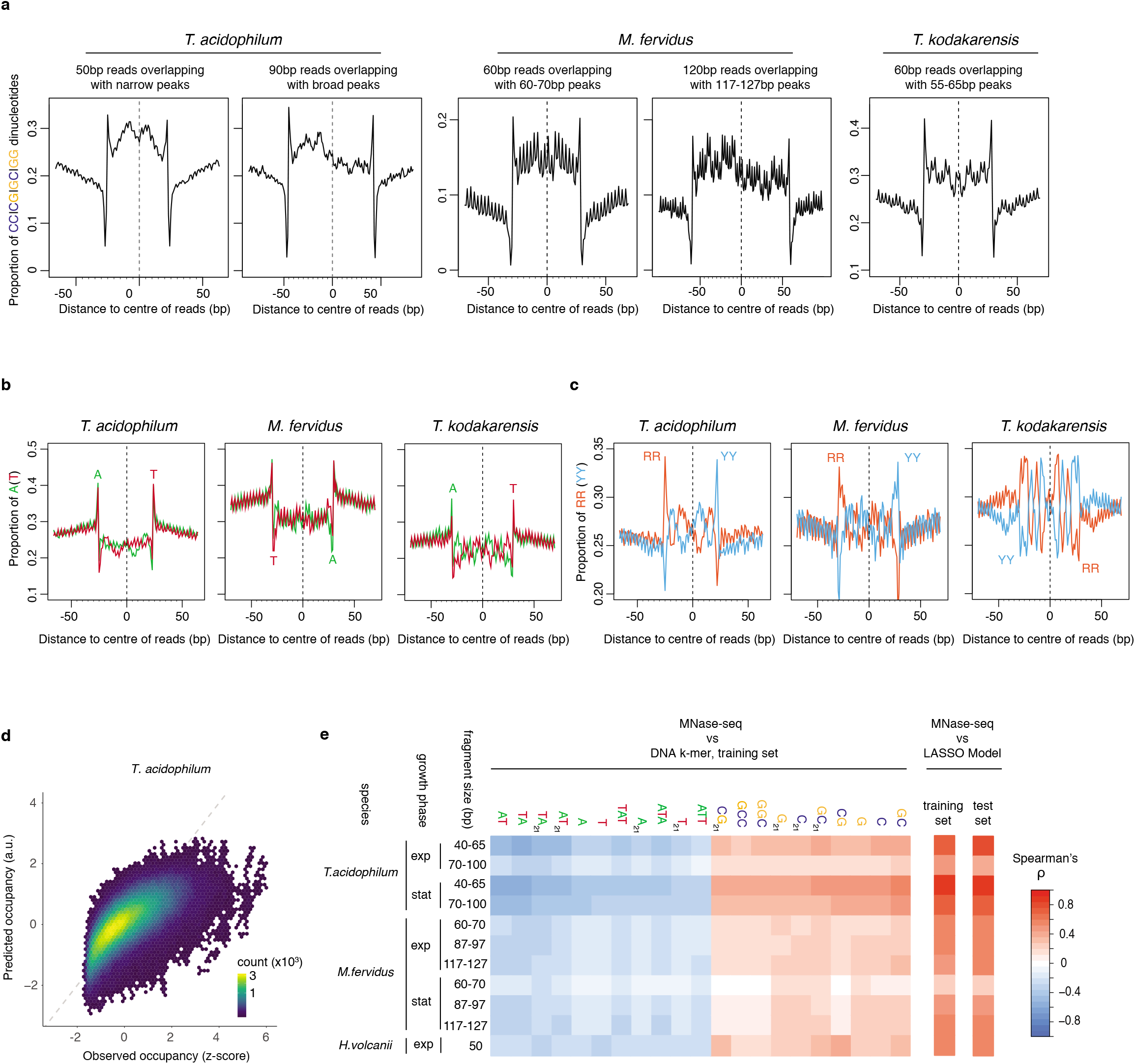
Comparison and predictive power of nucleotide enrichment patterns associated with HTa and archaeal histones. **(a)** Proportion of SS (= CCICGIGCIGG) dinucleotides, **(b)** AIT mononucleotides, and **(c)** RR (= purine/purine)lYY (= pyrimidine/pyrimidine) dinucleotides relative to the centres of reads of defined length in different archaeal species (see Methods for read filtering). **(d)** Density plot comparing observed (day 2, replicate 2) and predicted MNase-Seq coverage across the part of the *T. acidophilum* chromosome not used for training. **(e)** Correlation between MNase-seq coverage and individual DNA k-mers with particularly high positive or negative correlation coefficients, as observed in the training data. Overall correlations between measured MNase-Seq coverage and coverage predicted by the LASSO model, for both trained and untrained data, are shown on the right-hand side.

To define nucleotide preferences more rigorously and enable prediction of relative occupancy from underlying nucleotide features, as previously done for eukaryotic histones (Tillo and Hughes 2009), we trained Lasso models on small and large fragments separately (see Methods). The model confirms a general enrichment for strong (S = G or C) over weak (W = A or T) nucleotides, with mono- and dinucleotide frequencies the strongest individual predictors (Figure 5e). For small fragments, predicted and observed occupancy are well correlated (ρ (day 2)=0.76; ρ(day 3.5)= 0.86, observed versus predicted in test set, P<2e-16) although the model fails to capture extreme occupancy values (Figure 5d). The HTa occupancy signal we observe is, in a further parallel to histones, almost certainly conflated by MNase digestion bias, which is characterized by preferential degradation of AT-rich sequences (Allan *et al*. 2012). As a consequence, it would be premature to draw quantitative conclusions about the *extent* to which HTa prefers GC-rich sequences. However, not only do we observe read-internal nucleotide enrichments, which are not expected to arise as a consequence of digestion bias and have also been found in eukaryotes using chemical mapping approaches (Brogaard *et al*. 2012), but we also readily recover AT-rich protected fragments if they exist *in vivo*, as described in the following section. This indicates that, as previously shown for eukaryotes (Allan *et al*. 2012), inferred HTa/histone binding preferences are conflated with but not simply a mirage of MNase cutting bias.

### A *hidden diversity of large fragments in exponential phase*

Although the Lasso model does an admirable job of predicting small fragment occupancy across the growth cycle, it curiously performs much worse when trying to predict large fragment coverage in exponential phase (day 2, Figure 5e, ρ=0.43, P< 2.2e-16). We reasoned that differences between broad peaks in exponential and stationary phase might be owing to changes in the relative abundance of qualitatively different protective binding events: one, where large HTa-protected footprints emerge from extension of smaller fragments, and another where these fragments represent something else; a different mode of HTa binding that is independent of prior sequence-driven nucleation, perhaps, or an altogether different protein complex that happens to protect a similar-sized piece of DNA. To explore this hypothesis, we divided broad peaks into deciles based on the relative coverage of small fragments at these peak. Doing so, we find that broad peaks where small fragments are rarest show strongly divergent sequence composition (Figure 6a). Curiously, while these divergent peaks are very common in exponential phase, they almost entirely disappear in stationary phase, which is instead dominated by broad peaks that conform to the nucleation and extension model, where small fragment abundance quantitatively predicts large fragment abundance (Figure 6b, Figure S6).

**Figure 6.**
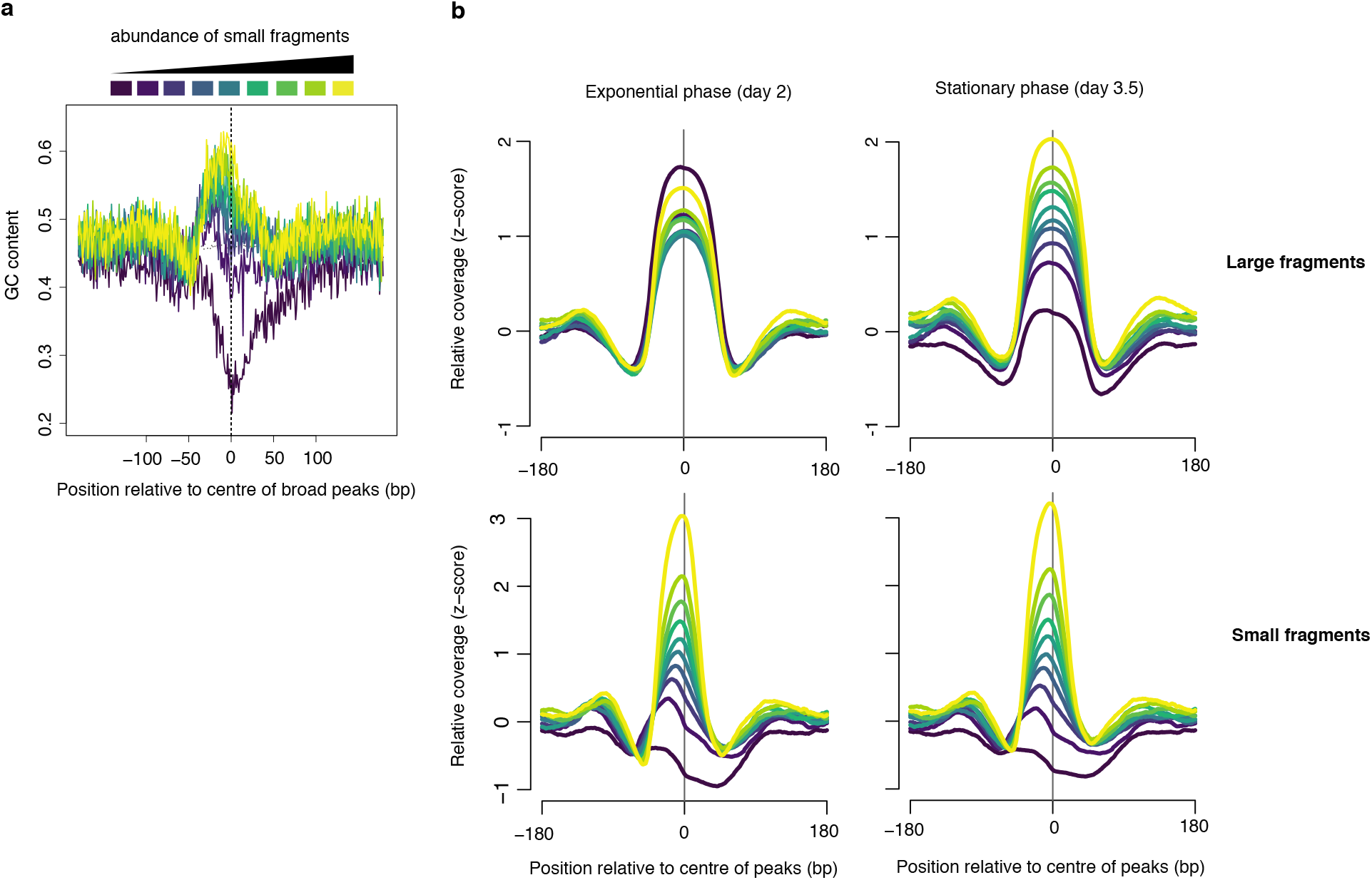
Broad peaks are associated with heterogeneous GC content in exponential but not stationary phase. **(a)** Average GC content at broad peaks (day 2), separated into deciles based on the relative abundance of small fragments and **(b)** the corresponding relative coverage for large and small fragments during exponential and stationary phase. For all graphs, decile decomposition is based on small fragment occupancy during exponential phase (day 2).

### Landscape of HTa binding around transcriptional start sites

Broad peaks with few small fragments are particularly enriched in intergenic sequence (Figure 7a), and it is around transcriptional start sites (TSSs, Figure 7b; approximated from RNA-seq data, see Methods) and end sites (TESs, Figure S7), where their disappearance in stationary phase is arguably most striking (Figure 7b). However, even though their positioning suggests a potential involvement in gene regulation, their disappearance is not obviously coupled to local changes in transcription: the abundance of these “independent” fragments drops equally for genes that are up- or down-regulated or remain the same in stationary compared to exponential phase (Figure 7f). We find no equivalent to these nucleation-independent large fragments in histone-encoding archaea (Figure 7c-e, Figure S7) and further investigation will be required to establish the identity of the proteins that protect these fragments from digestion. We note, however, that nucleoid-associated proteins in bacteria, including HU and H-NS in *E. coli*, can bind different DNA substrates in different conformations and therefore exert distinct effects on DNA topology and chromatin architecture depending on local concentration, sequence context, and the presence of other factors (Dillon and Dorman 2010). It is therefore tempting to speculate that HTa has retained some of this HU-like versatility in DNA binding and that independent fragments might also be derived from HTa protection. However, we have no direct evidence for this at present. Small fragment-peaks and nucleation-dependent large fragments, on the other hand, share substantial similarities with archaeal histones in their positioning around genes: they are depleted at all times from the AT-rich TSSs/TESs, and barely change position through the growth cycle (Figure 7, Figure S8).

**Figure 7.**
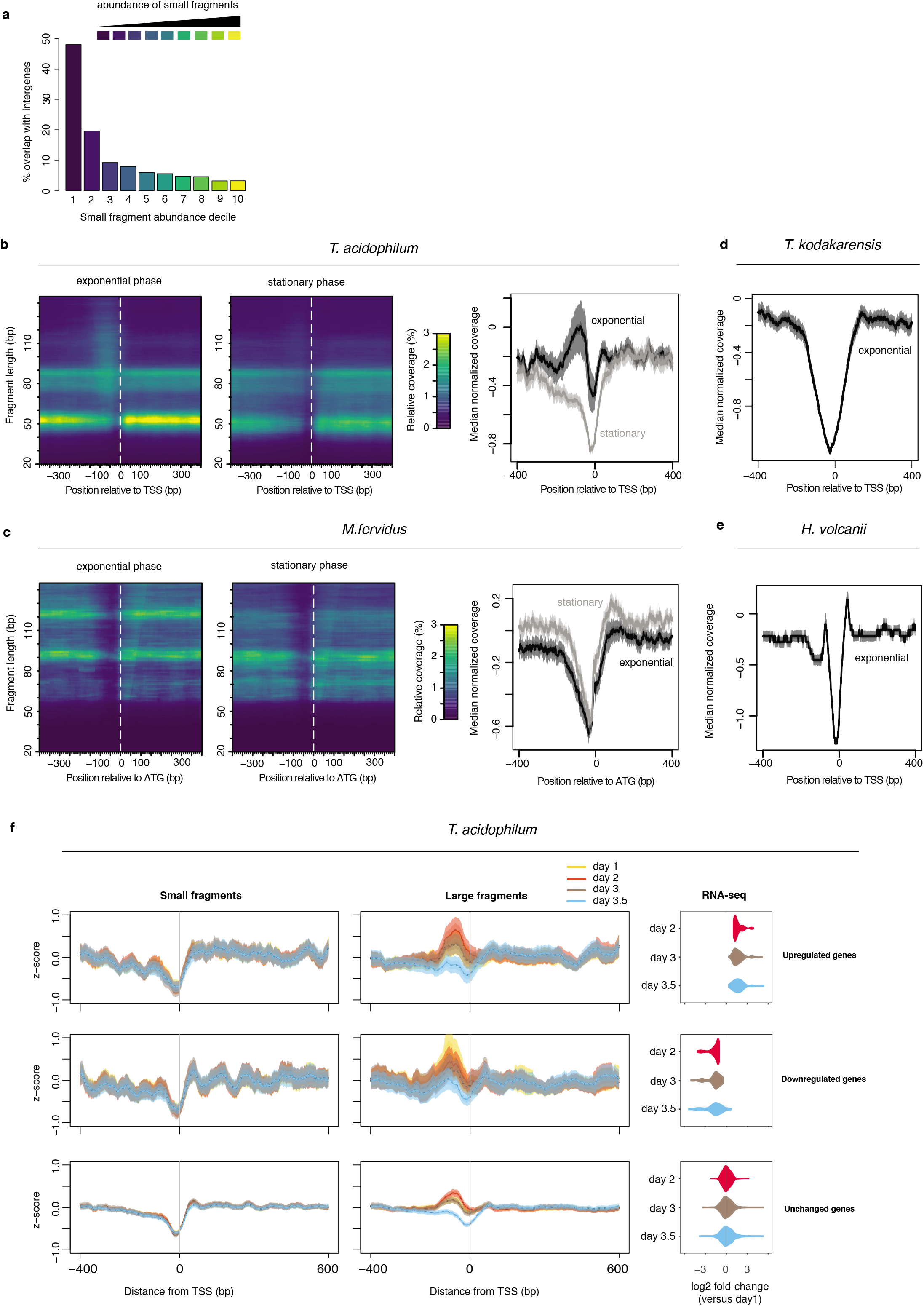
MNase-Seq coverage around transcriptional start sites in *T. acidophilum* and histone-encoding archaea in the context of dynamic transcription. **(a)** Broad peaks associated with low abundance of small fragments are enriched in intergenic regions. **(b)** Left and central panel: Heat maps indicating MNase-seq coverage by fragment length relative to transcriptional start sites in exponential (day 2) and stationary phase (day 3.5). Right panel: median normalized MNase-seq coverage (considering all fragment sizes) as a function of distance from the TSS. **(c)** as in (b) but for *M. fervidus* and using the coding start (ATG) rather than the TSS as a reference point. To ensure that the coding start constitutes a reasonable proxy for the TSS, only genes with a divergently oriented neighbouring gene are considered, thus eliminating genes internal to operons. **(d, e)** median of normalized MNase-seq coverage (considering all fragment sizes) as a function of distance from the TSS in *T. kodakarensis* and *H. volcanii*. **(f)** Changes in normalized MNase-seq coverage for small and large fragments around transcriptional start sites in *T. acidophilum* as a function of growth phase and whether genes are upregulated, downregulated or remain unchanged relative to RNA abundance on day 1. Genes are grouped according to differential expression (or lack thereof) on day 2 compared to day 1. Genes with a log2-fold change >1 were considered significantly upregulated, those with a log2-fold change <−1 significantly down-regulated (FDR<0.01). The rightmost panels indicate that a majority of genes up-/downregulated on day 2, remain up-/downregulated on days 3 and 3.5.

## Discussion

The evidence above suggests that HTa is a protein of bacterial origin that converged onto supramolecular properties reminiscent of archaeal histones. Whereas well-characterized HU homologs exhibit elevated occupancy in AT-rich regions, prefer an AT-rich motif at the centre of the binding footprint, or bind sequence non-selectively (Swinger and Rice 2007; Grove 2011; Prieto *et al*. 2012), HTa shows a more histone-like preference for GC and even exhibits asymmetric mononucleotide/RR/YY profiles across the dyad. Whether these shared nucleotide preferences are grounded in primary sequence, DNA structural features determined by primary sequence, or a combination of the two remains to be established. Sequence-driven positioning around functional genomic landmarks, footprint extension dynamics, and relative positional stability throughout the growth cycle are also reminiscent of archaeal histones, as is the ability of HTa to protect DNA from MNase digestion. Based on these observations, we propose that HTa can be regarded as an archaeal histone analog.

This analogy is, of course, preliminary. Structural work will be required to characterize how HTa interacts with DNA and to determine whether large HTa-protected fragments reflect closely spaced binding events or, in further analogy to archaeal histones, the presence of contiguous HTa polymers. The observation that even larger protected fragments can be formed when HTa is expressed in *E. coli* (Figure S1a) is interesting in this regard and deserves further exploration. Additional work is also required to elucidate the interaction partners and physiological functions of HTa, including its involvement in transcription. Although we find no obvious global link between changes in HTa occupancy and transcriptional output, this does not preclude local effects or dynamics at a time-scale where information on nascent transcription – rather than steady-state RNA levels – would be required to implicate HTa. Neither do our results rule out an important but constitutive role in transcription, for example in binding to and constraining negative supercoils similar to what has been observed for other HU homologs (Berger *et al*. 2009; Grove 2011) and archaeal chromatin proteins such as MC1, Sul7, and Cren7 (Zhang *et al*. 2012). Alternatively, the main function(s) of HTa might simply not be in transcription. Searcy and co-workers showed that HTa facilitates re-annealing following DNA denaturation (Stein and Searcy 1978), suggesting that HTa might assist DNA renaturation after thermal stress. Our finding of heterogeneous but predictable HTa occupancy across the genome implies that re-annealing would proceed differentially in genic and intergenic regions, leaving the latter unbound and free to engage in binding of protein complexes for transcription. However, the physiological importance of HTa-mediated reannealing remains to be established and is put into perspective by the observation that *E. coli* HU, when added to naked DNA, also significantly raises its melting temperature (Drlica and Rouvière-Yaniv 1987). Another possibility is that HTa is involved in higher order genome structure (Bohrmann *et al*. 1990) or, like many HU family members (Grove 2011), in DNA repair. To dissect these and other hypothetical functions *in vivo* in the future, to determine whether HTa is essential, and address whether archaeal histones can functionally substitute for HTa, the development of genetic tools for *T. acidophilum* is highly desirable.

Finally, from an evolutionary perspective, it is worth highlighting dinoflagellates as a second case where histones have lost their pre-eminent role in genome compaction and organization to other small basic proteins. Even though histones remain encoded in dinoflagellate genomes (Marinov and Lynch 2016), and might continue to play important roles in transcription or other processes, they are no longer the main protein constituent of chromatin. Instead, proteins with homologs in phycodnaviruses have become the principal mediators of compaction (Gornik *et al*. 2012). In a subset of species, these dinoflagellate/viral nucleoproteins (DVNPs) act alongside HU-like proteins that were likely acquired from bacteria (Figure 2) (Wong *et al*. 2003; Janouškovec *et al*. 2017) and might be involved in the genesis and maintenance of DNA loop structures (Chan and Wong 2007). If not exactly a precedent, the case of dinoflagellates nonetheless serves to illustrate that HU proteins are versatile, evolvable and have been independently co-opted into important architectural roles following horizontal transfer. It also highlights that histones or a subset of their functions can be replaced, under the right circumstances, by alternative DNA-binding proteins. At the same time, our results demonstrate that such replacements, even though they appear radical, need not necessarily go hand in hand with fundamental changes to the architectural layout of chromatin.

## Methods

### Thermoplasma acidophilum *culture*

*T. acidophilum* strain 122-1B2 was obtained from DSMZ (https://www.dsmz.de/) and cultured using the medium described in (Searcy and Stein 1980), supplemented to a final concentration of 2g/L yeast extract (BD Biosciences). The medium was boiled for five minutes and allowed to cool to 58°C before inoculation. Cultures were incubated at 59°C with shaking (90rpm) in an INFORS Thermotron incubator. Throughout, culture to flask volume ratio was maintained at a maximum of one fifth. Fresh cultures were inoculated with a 10% v/v inoculum from a 4-day-old culture. Samples from 24-, 48-, 72-, and 80-hour cultures were used for RNA extraction and MNase experiments. Sample aliquots were first equilibrated to pH4 using NH_4_OH to avoid depurination (Robb *et al*. 1995) and then pelleted at low speed. To monitor cell viability and homogeneity, cells were imaged with a standard light microscope at 100x magnification.

### *MNase digestion* – Thermoplasma acidophilum

*T. acidophilum* cells self-lyse at pH>6 (Robb *et al*. 1995). Pellets were therefore directly re-suspended in ice-cold MNase digestion buffer (10mM Tris, 5mM Ca^2+^, pH8) and homogenized via 20 passages through a Dounce homogenizer, on ice. Unfixed crude lysates were digested in the presence of 4U/mL of MNase (Thermo Fisher) at 37°C. Digestion was stopped by addition of EDTA to a final concentration of 20mM. Samples were incubated for an additional 30min at 37°C in the presence of RNAse A to a final concentration of 0.5mg/mL and then overnight at 65°C in the presence of SDS (1%) and proteinase K (125μg/mL). Subsequently, DNA was extracted by phenol chloroform extraction and precipitated with ethanol. For MNase digestion of *E. coli* samples, lysates were obtained by cryo-grinding with a pestle and mortar.

### Preparation of undigested DNA samples

Undigested lysate was incubated as above but without addition of enzyme. Undigested DNA was then sonicated using a Covaris S220 sonicator with the following settings: peak power: 175, duty: 10, cycle/burst: 200 for 430s (target size: 150bp).

### *MNase digestion and sequencing* – Methanothermus fervidus

Flash-frozen pellets of *M. fervidus* harvested in late exponential and stationary phase were purchased from the Archaeenzentrum in Regensburg (via Harald Huber). Digestion and sequencing conditions will be published as part of a manuscript currently in preparation. They are not repeated here but reproduced in full at the NCBI Gene Epression Omnibus (GEO) where the raw reads have been deposited (accession number pending).

### RNA extraction

Pellets were re-suspended in 500*μ*L RNAlater (Ambion), incubated for one hour at room temperature, pelleted again and snap-frozen. Total RNA was extracted using an RNeasy kit (Qiagen) according to manufacturer’s instruction, including DNAse I treatment.

### Protein expression in E. coli

Protein sequences for HTa and different HU homologs were obtained from UniProt (P02345, P0ACF0, LOEKC1, A0A0R2DT96). The corresponding coding sequences were codon-optimized for expression in *E. coli*, synthesized and inserted into the pD864-SR backbone for rhamnose-inducible expression (ATUM). The YFP-expressing control strain was obtained directly from ATUM.

### Bacterial strains and growth conditions

*E. coli* ΔΔhupA/hupB double mutant and corresponding wildtype strain were a gift from Jacques Oberto. These strains were grown in LB medium at 37°C with shaking. For MNase assays, strains were pre-cultured from selection plates to 5mL LB cultures in 50mL falcon tubes overnight with antibiotics and re-suspended to 0D600=0.1. 1M rhamnose was added two hours after inoculation to a final concentration of 5mM and grown for 4 hours prior to harvesting. Induction was confirmed by monitoring YFP expression.

### Sequencing and data availability

For MNase digestion experiments paired-end reads were prepared using the NEBNext Ultra II DNA Library Prep Kit. For RNA sequencing, ribosomal RNAs were depleted using Illumina’s Ribo-Zero rRNA Removal Kit (Bacteria) and RNA libraries prepared using the TruSeq Stranded Total RNA LT Kit. Both RNA and DNA libraries were then sequenced on a HiSeq2500 machine. Raw read and processed data have been deposited in GEO (accession number pending).

### MNase-seq analysis

Paired-end 100bp reads were first trimmed using BBDuk v37.64 (parameters: ktrim=r, k=21, hdist=1, edist=0, mink=11, qtrim=rl, trimq=30, minlength=10, ordered=t, qin=33) and merged using BBmerge v37.64 (parameters: mininsert=10, mininsert0=10). The merged reads were then mapped to the *T. acidophilum* DSM1728 genome (GCA_000195915.1) using Bowtie2 (-U). Coverage tracks were computed using bedtools bamCoverage (RPGC normalization, effective genome size: 1564906bp), measuring coverage for reads of sizes 40-65bp and 70-100bp separately where appropriate. To compute the correlation matrix of reads of different sizes (Figure 3d), coverage for reads of defined lengths was computed using bedtools bamCoverage without RPGC normalization.

### Normalization for replication associated bias

Reads from the sonicated DNA control samples were mapped in paired-end mode using Bowtie2. Coverage was computed independently of read size and smoothed over 10kbp to avoid introducing noise from the input into the MNase signal. Genome-wide coverage of MNase-digested samples was then divided by its corresponding undigested DNA sample to remove bias in coverage associated with replication. Lastly, coverage was converted into a Z-score using the *scale* function in R.

### RNA-seq analysis

Single-end reads were mapped to the *T. acidophilum* DSM1728 genome using Geneious 11.1.2. Transcripts were reconstructed and quantified using Rockhopper (McClure *et al*. 2013), using a stringent threshold of 0.9 for 5’ and 3’UTR detection. Differential expression was assessed using DESeq2 as part of the Rockhopper pipeline.

### LASSO modeling

Abundances of different nucleotide k-mers k={1,2,3,4} were computed over genomic windows of 21bp and 51bp using the R packages seqtools (v1.16). We included the 21bp window to enable independent capture of read-internal nucleotide patterns that did not overlap with MNase cutting sites. For all genomes analyzed, the top 80 k-mers (across both 21bp and 51bp windows) with the highest absolute correlation values with MNase coverage over the first third of the genome were chosen as model parameters. A general linear model with LASSO optimization was trained on the first third of each genome, with 10-fold internal cross validation, using the LASSO function in Matlab. Predicted coverage tracks were then calculated in R based on k-mer weights from the training set.

### Peak detection and asymmetry scoring

Peaks were detected using the NucleR R package and scored using a modified scoring function, where peak height was measured as the coverage value at each peak centre relative to the empirical distribution of the data. Parameters used for the initial Fourier filtering step are listed in Table S3. A threshold value corresponding to a Z-score of 0.25 was used for all data. To score asymmetric coverage inside broad peaks, we computed the average coverage signal of smaller DNA fragments either side (*a* and *b* in Figure 4b) of the peak dyad and then considered the ratio *a*/*b*. Coverage signal at peaks where *a*/*b*<1 were flipped around the dyad axis for downstream signal processing.

### Nucleotide frequencies and data visualization

Nucleotide frequencies were computed using the R package Seqpattern (v1.14). Only reads with an average quality score of 30 that overlapped called peaks by ≥90% were selected for this analysis. 2D occupancy plots were generated from non-normalized MNase-Seq reads using plot2DO (https://github.com/rchereji/plot2DO) (Chereji *et al*. 2018), modified to enable processing of single-end reads. Multiscale analysis of MNase coverage was carried out using MultiScale Representation of Genomic Signals (https://github.com/tknijnen/msr/) (Knijnenburg *et al*. 2014), considering genomic segments with significant (P<1e-10) enrichment at scale 30.

### MNase and additional data from other organisms

MNase data from other organisms was obtained from the Sequence Read Archive (SRR495445 for *T. kodakarensis*; SRR574592 for *H. volcanii*). *H. volcanii* reads were trimmed using BBDUK (ktrim=r, k=21, hdist=1, edist=0, mink=11, qtrim=rl, trimq=20, minlength=10, qin=33). Reads were mapped to their respective genomes using Bowtie2 (Setting for SRR495445 reads: -3 75 -X 5000 -k 1 -x, as in the original publication; default settings for SRR574592 reads). To reduce PCR duplicate bias, per base coverage values of MNase data from *H.volcanii* were thresholded at the last percentile. TSS positions for *T. kodakarensis* and *H. volcanii* were obtained from (Jäger *et al*. 2014) and (Babski *et al*. 2016), respectively.

### Homology modelling

Secondary structures for HupA (P0ACF0) and HmfA (P48781), as displayed schematically in Figure 1a, were taken from UniProt. The secondary structure of HTa was predicted by homology modeling in SWISS-MODEL (Waterhouse *et al*. 2018) using HupA (PDB:1mul) as a template. To predict and visualize the quaternary structure of the HTa homodimer, we used the HTa sequence to build a homology model based on the X-ray crystal structure of (HupA)_2_ using PDB: 1p51 as a template. The homology model, again built using SWISS-MODEL, has a general mean quality estimate of 0.71. Both (HTa)_2_ and (HupA)_2_ structures were refined with steepest descent and conjugate gradient energy minimization using the AMBER ff14SB protein force field potentials (Maier *et al*. 2015) and a force constraint of 2 kcal/mol placed on the Ca peptide backbone atoms. To calculate the solvent accessible surface area and charge density, we used the Adaptive Poisson-Boltzmann Solver (APBS) (Jurrus *et al*. 2018). The charge density was mapped onto the solvent accessible surface area using the VMD visualisation package (Humphrey *et al*. 1996).

### Phylogenetic analysis

Amino acid sequences containing the HU-IHF domain (cl00257) where identified in bacteria, archaea, eukaryotes, and viruses using the Conserved Domain Architecture Retrieval Tool (Geer *et al*. 2002) [accessed on 29th October 2018]. The initial set comprised 52,953, 82, 204, and 131 sequences, respectively. To reduce the number of bacterial sequences to a computationally more tractable subset yet maintain sequence diversity, we pre-processed the bacterial sequence set as follows: first, each bacterial sequence was identified to family level using NCBI taxonomy annotations; then, only those sequences with a valid family-level taxonomic identification were retained (43,454 sequences belonging to 409 families). Within each family, we then calculated pairwise identities between all sequences and identified up to ten sequence identity clusters. Subsequently, a single representative sequence was randomly sampled from each cluster (for families comprising of fewer than 10 sequences, we selected all available sequences). This reduced the bacterial set to 3135 sequences.

Next, the full archaeal, eukaryotic, viral and reduced bacterial sets were processed using the Batch Web CD-Search Tool (Marchler-Bauer and Bryant 2004) to determine the position and integrity of the HU-IHF domain(s). Based on the domain identification, we then selected only those sequences from each set that contained a single, complete domain. The bacterial set was further reduced by selecting only those sequences less than or equal to 110 amino acids. Given the relative scarcity of sequences in the other kingdoms, their sequences were not size-filtered. The final set comprised 30 archaeal, 164 eukaryotic, 112 viral, and 1920 bacterial sequences.

Sequences were aligned using the Constraint-based Multiple Alignment Tool (COBALT) through the NCBI web-interface (https://www.ncbi.nlm.nih.gov/tools/cobalt/cobalt.cgi) with default parameters. The phylogenetic tree was reconstructed using RAxML (version 8.2.10) with the following parameters: -f a, -m PROTCATAUTO, -T 20. Branch support was based on 100 bootstrap calculations performed in RAxML (Stamatakis 2014).

## Acknowledgements

The authors thank Dennis Searcy for advice on *T. acidophilum* husbandry and chromatin biochemistry, Finn Werner for mentorship and his lab for feedback and advice, Harald Huber and the Archaeenzentrum in Regensburg for *M. fervidus* biomass, and Jacques Oberto for *E. coli hupA/B* deletion strains. This work was supported by a UKRI Innovation Fellowship (JBS), an Imperial College Integrative Experimental and Computational Biology Studentship (AE), and UK Medical Research Council core funding (TW).

## Author contributions

AH carried out all experiments and analyses, with the following exceptions: MR produced and contributed to analysis of *M. fervidus* MNase data; JBS carried out homology modelling; and AE designed and implemented pipelines for HU ortholog identification and phylogenetic analysis. TW conceived the study. AH and TW devised experimental and analytical strategies and wrote the manuscript, with input from all authors.

