## Supplementary Material for "The DNA-binding protein HTa from *Thermoplasma acidophilum* is an archaeal histone analog"

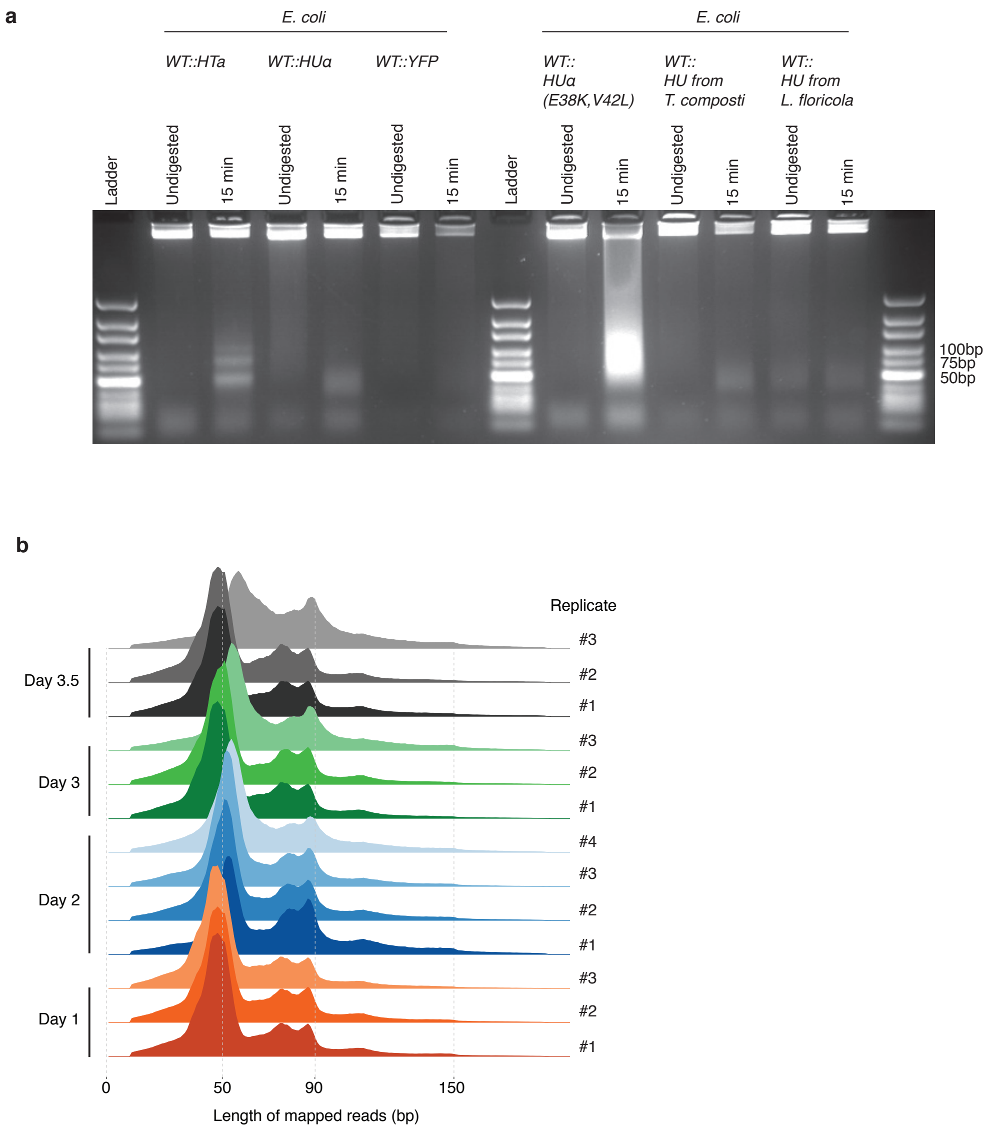


**Figure S1. (a)** Agarose gel (3%) of MNase digestion products from *T. acidophilum* (day 2) along with digestion products of *E. coli* ectopically expressing either HTa, HupA, YFP, HupA (E38K,V42L), HU from *T. composti* or HU from *L. floricola*, from the same plasmid backbone. HupA (E38K,V42L) is a mutant that had previously been shown to induce extreme compaction of the *E. coli* nucleoid (Kar et al. 2005 PNAS 102:16397-16402). **(b)** Distribution of the lengths of fragments mapped to the *T. acidophilum* genome for all replicates across the growth cycle.

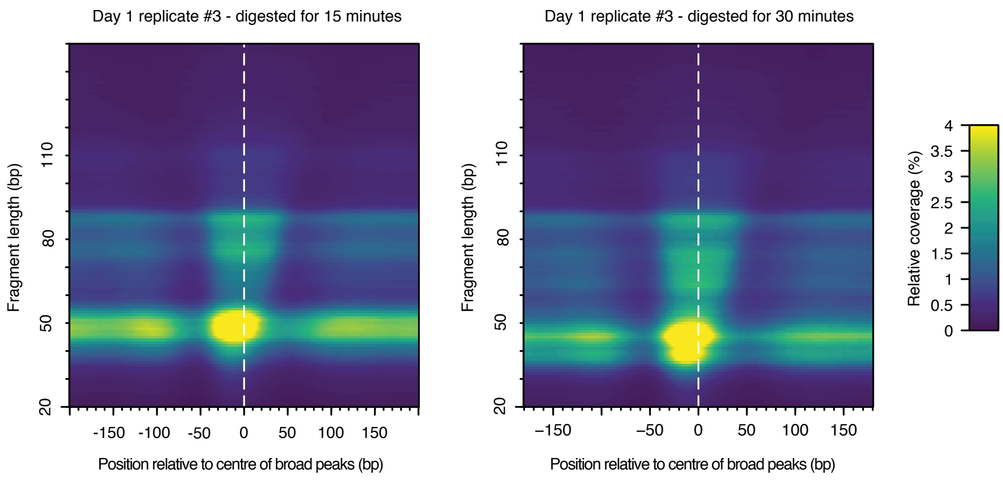


**Figure S2.** Heat maps indicating MNase-seq coverage by fragment length relative to the centre of broad peaks in *T. acidophilum*, for the same sample (day1, replicate 3), digested for either 15 or 30 minutes.

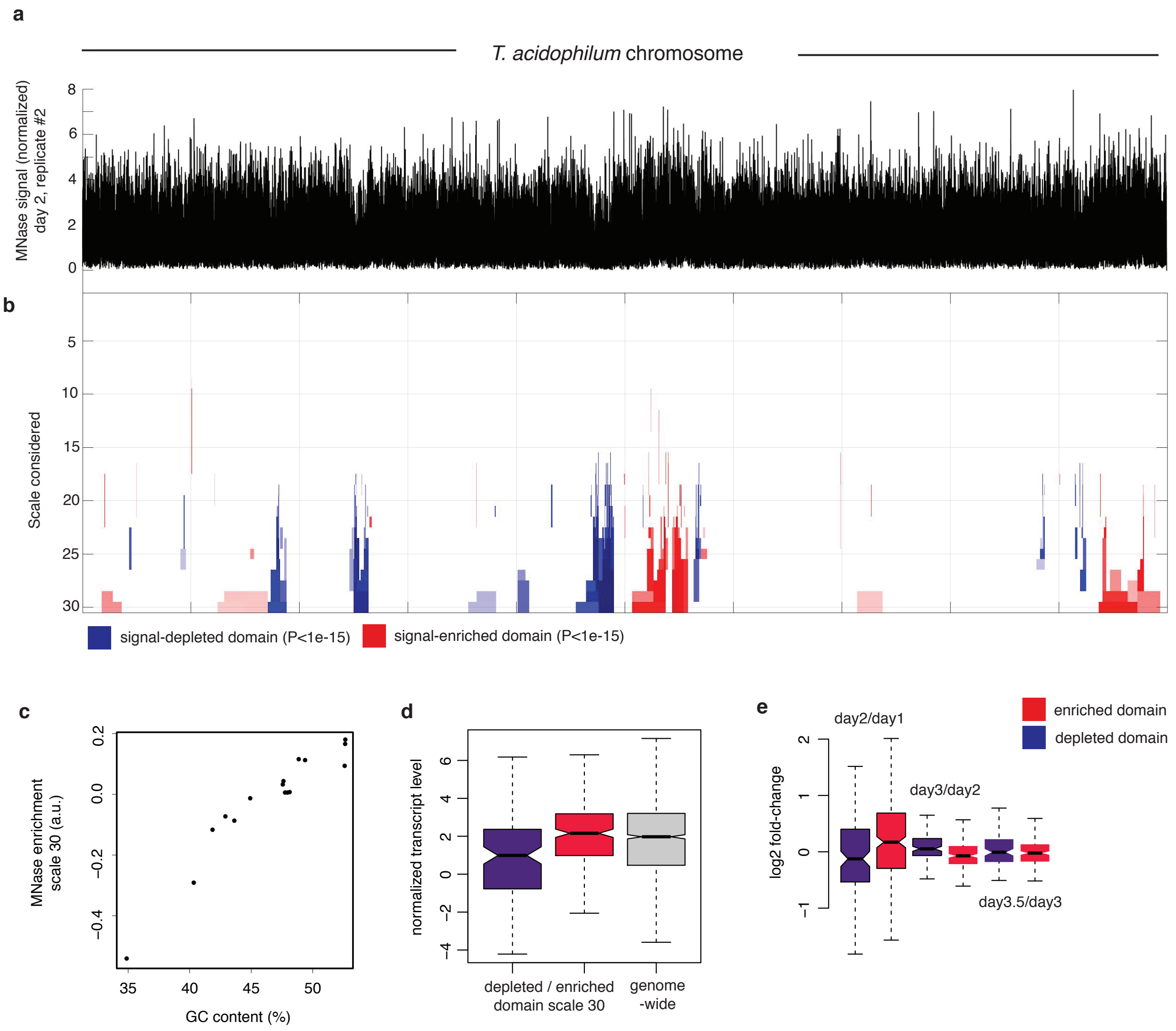


**Figure S3. (a)** Chromosome-wide normalized MNase-Seq coverage along the *T. acidophilum* chromosome (day2, replicate 2). **(b)** Multiscale analysis of MNase signal enrichment. Significantly enriched or depleted (p-value < 1.e-15) segments are colour-coded red and blue, respectively. Scales correspond to increasing window sizes over which enrichment is computed. **(c)** Enrichment signal of significantly MNase-signal-enriched or -depleted genomic domains at scale 30 as a function GC content. **(d)** Normalized transcript levels for pooled depleted or enriched domains at scale 30 and **(e)** corresponding log2-fold changes in transcript levels.

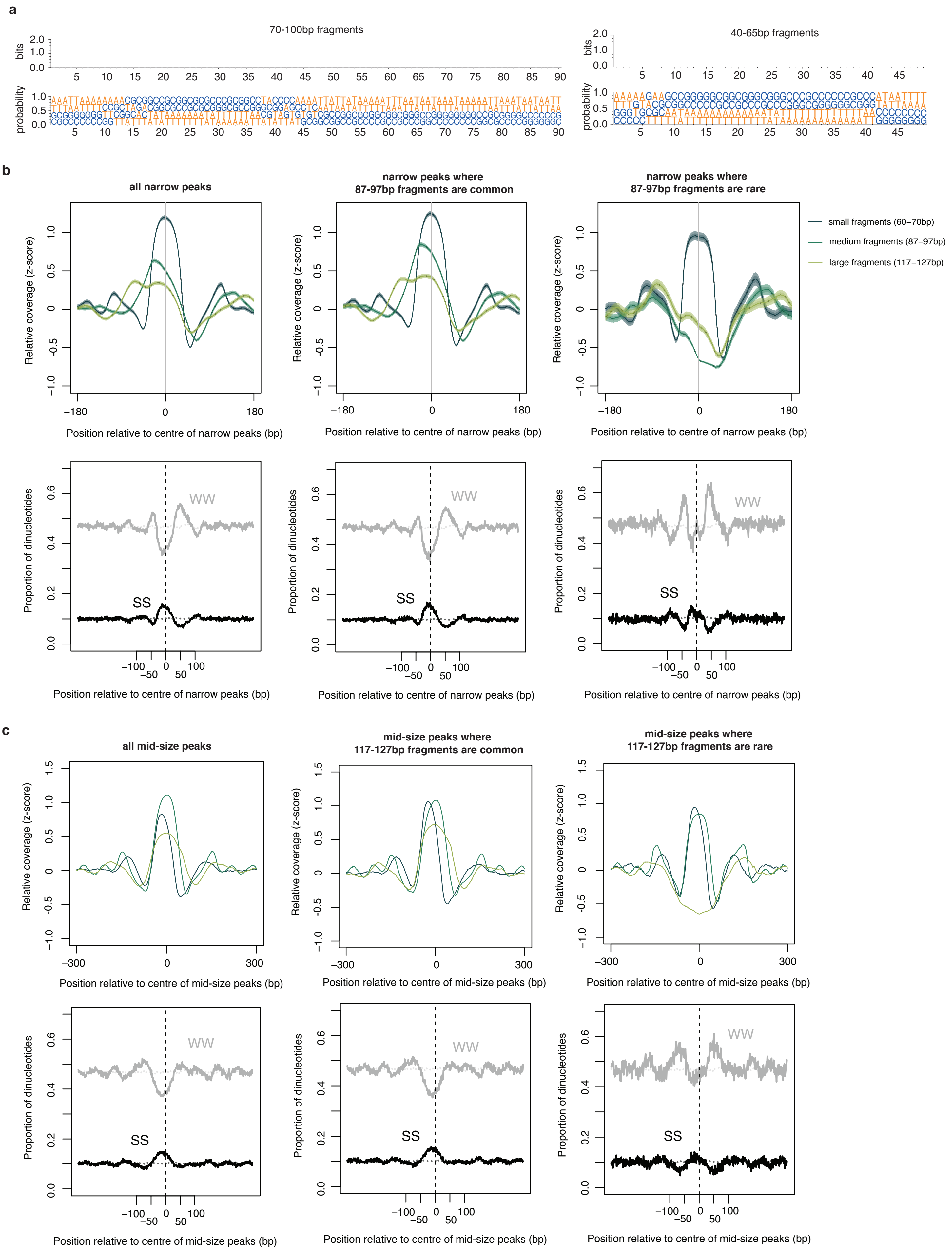


**Figure S4. (a)** Weblogos (upper panels) and nucleotide occurrence probabilities (lower panels) at narrow and broad peaks detected during exponential phase in *T. acidophilum*. Information content is so low that, when using the common 0-2 bit visualization range, the plots appear empty. (b) Normalized MNase-Seq coverage relative to the centre of narrow peaks oriented according to the abundance of 87-97bp fragments in *M. fervidus*. Middle and right panels are focused on peaks where 87-97bp fragments are common or rare, respectively. Lower panels display the proportion of SS (=CC|CG|GC|GG) and WW (=AA|AT|TA|TT) dinucleotides at locations matching the upper panels. Dotted lines indicate the proportion of SS or WW dinucleotides expected by chance, estimated by randomly sampling 25000 regions per genome. (c) as in (b) but for 87-97bp peaks scored according to 117-127bp fragments and oriented according to 60-70bp fragments. Note the increase in WW content flanking the smaller-sized peaks that do not get extended further.

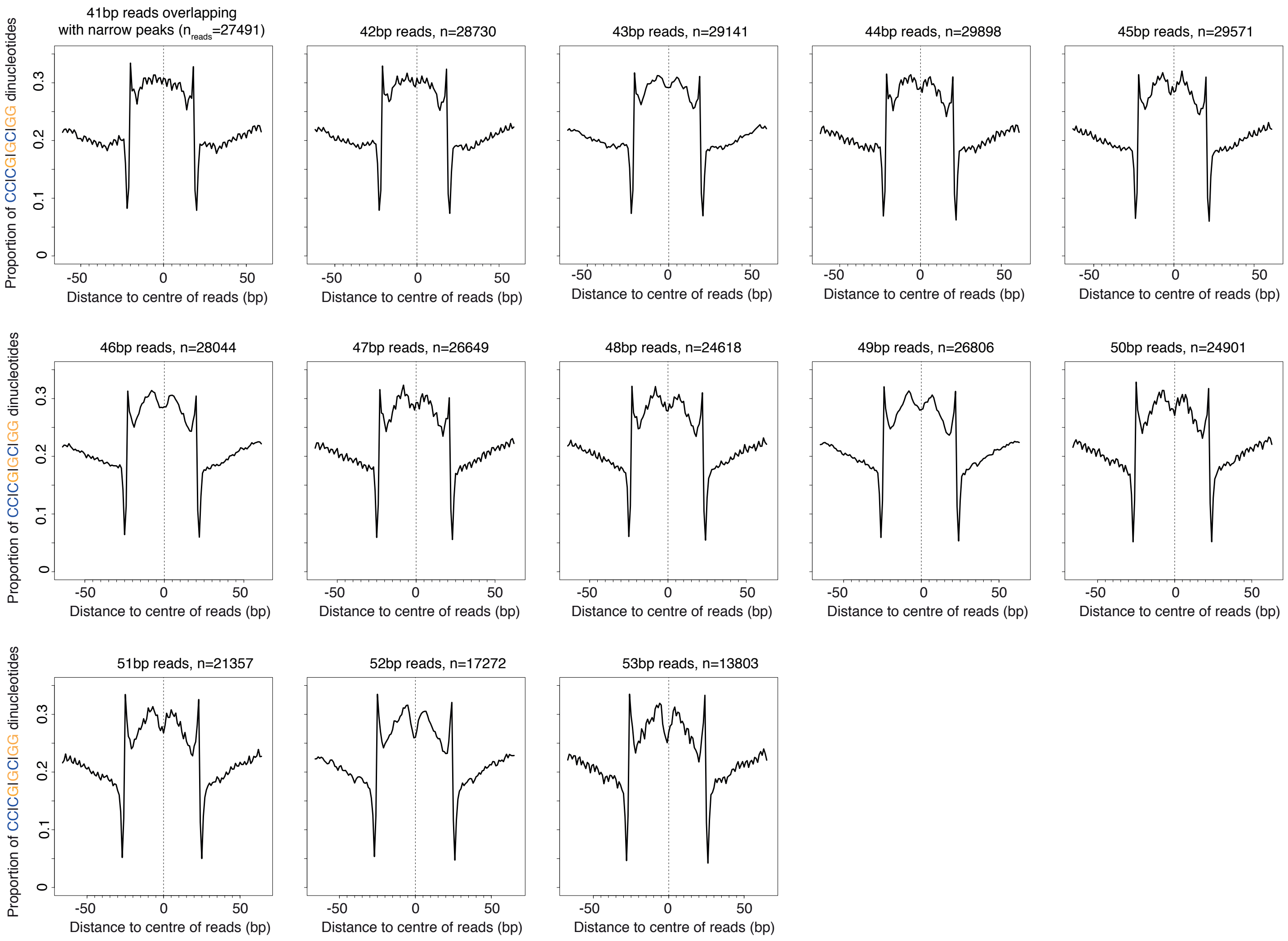


**Figure S5.** Proportion of SS (= CC|CG|GC|GG) dinucleotides relative to the centres of reads of defined length (41-53bp) in *T. acidophilum*.

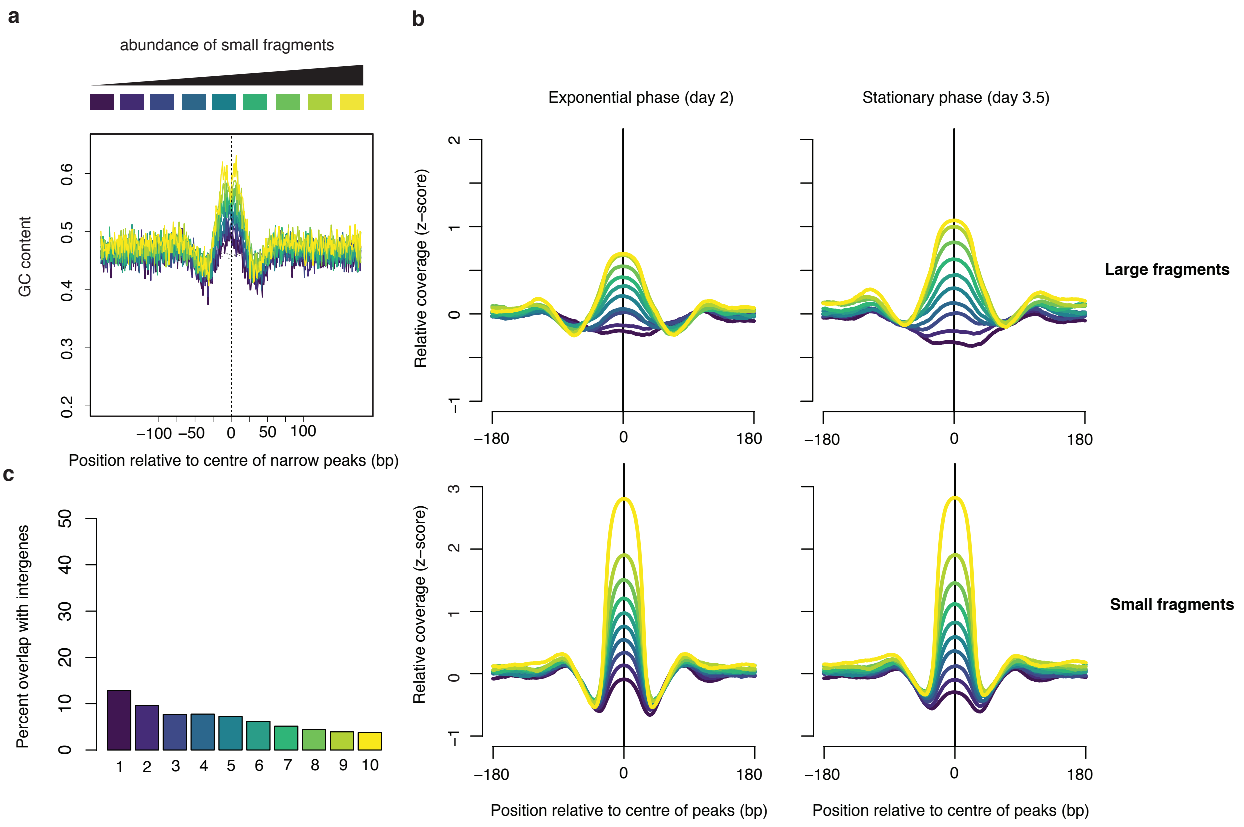


**Figure S6. (a)** Average GC content at narrow peaks (day 2), separated into deciles based on the relative abundance of small fragments. **(b)** corresponding relative coverage for large and small fragments during exponential and stationary phase. **(c)** Percentage of overlap between narrow peaks and intergenic regions. For all graphs, decile decomposition is based on small fragment occupancy during exponential phase (day 2).

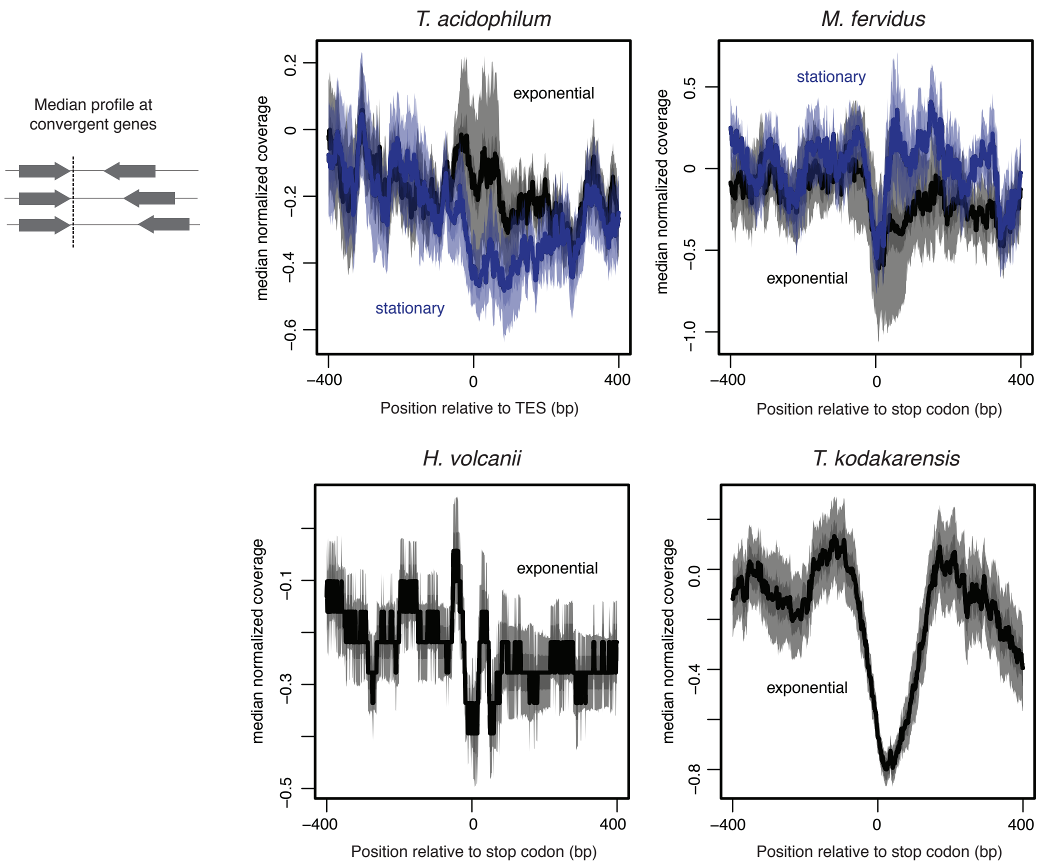


**Figure S7.** Median normalized MNase-seq coverage across fragment sizes relative to the distance from TESs or stop codons in different species. To ensure that the stop codons constitute a reasonable proxy for the TES, only genes with a convergently oriented downstream neighbouring gene are considered, thus eliminating genes internal to operons.

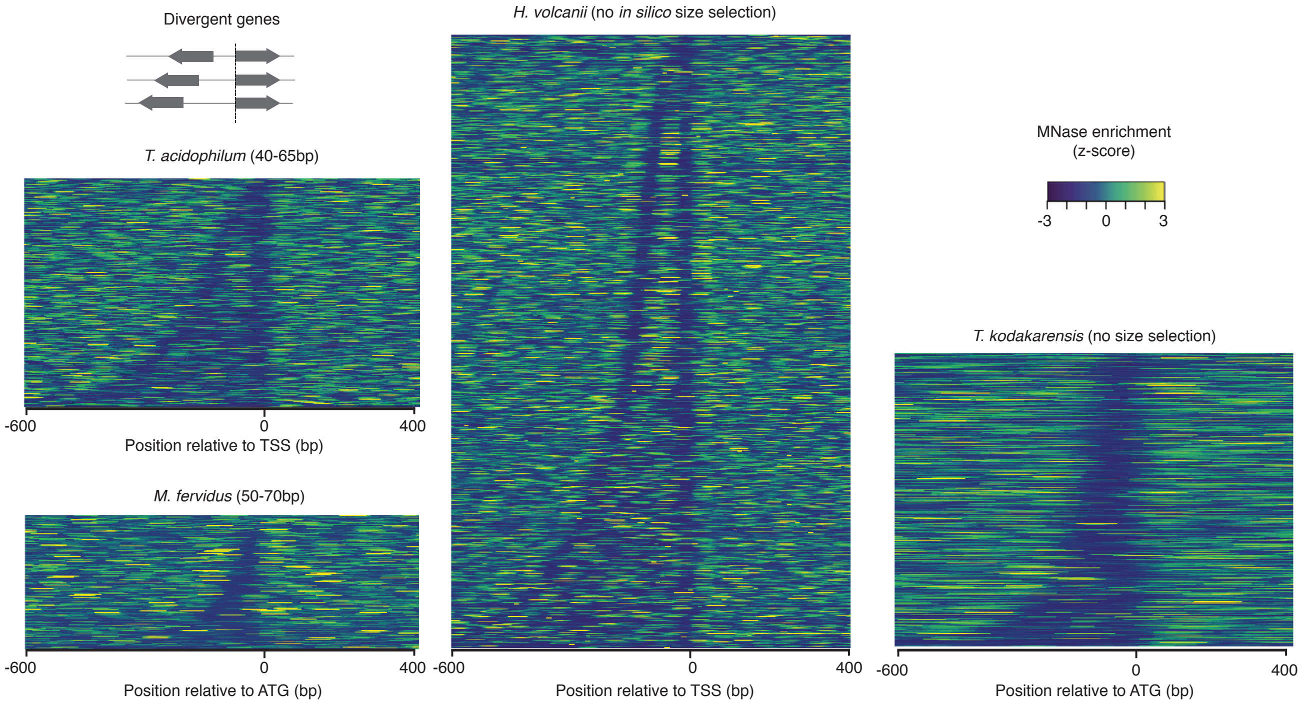


**Figure S8.** Heat maps displaying normalized MNase-seq coverage at divergent genes relative to the distance from the start codon (ATG) or TSS in different species. Intergenic regions are sorted according to their width.

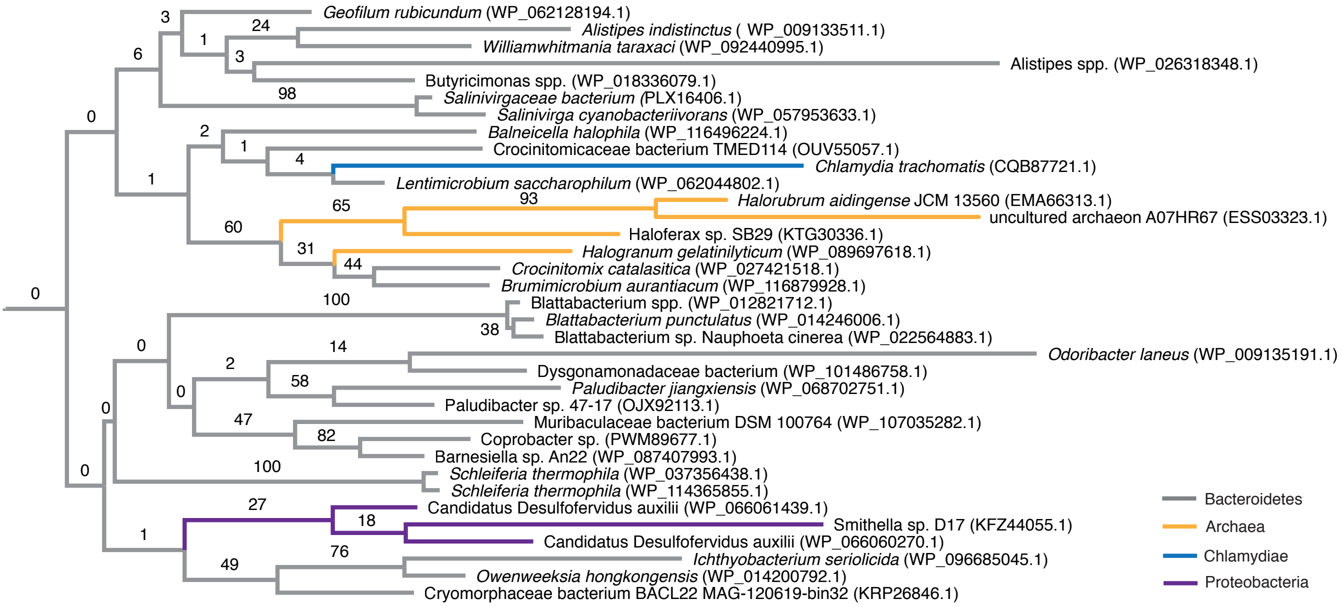


**Figure S9. Phylogenetic placement of HU proteins attributed to halophilic archaea.**
The phylogenetic tree shown is an excerpt of the protein-level HU family tree shown in Figure 2, focussing on sequences from halophilic archaea (orange), which cluster mainly with sequences of bacteria from the phylum Bacteroidetes (grey). As is true for the majority of the HU protein tree, deeper ancestral relationships are poorly resolved.

**Table S1. Representation of HU homologs across bacterial phyla.**

| **Phylum** | **Number of homologs detected** |
|---|---|
| Acidobacteria | 22 |
| Actinobacteria | 132 |
| Aquificae | 25 |
| Bacteroidetes | 142 |
| Caldiserica | 4 |
| Chlamydiae | 16 |
| Chlorobi | 4 |
| Chloroflexi | 25 |
| Chrysiogenetes | 4 |
| Cyanobacteria | 153 |
| Deferribacteres | 10 |
| Deinococcus–Thermus | 14 |
| Dictyoglomi | 2 |
| Elusimicrobia | 9 |
| Fibrobacteres | 8 |
| Firmicutes | 273 |
| Fusobacteria | 13 |
| Gemmatimonadetes | 10 |
| Lentisphaerae | 2 |
| Nitrospirae | 4 |
| Planctomycetes | 14 |
| Proteobacteria | 866 |
| Spirochaetes | 22 |
| Synergistetes | 7 |
| Tenericutes | 15 |
| Thermodesulfobacteria | 8 |
| Thermotogae | 35 |
| Verrucomicrobia | 47 |

**Table S2. Examples of putative archaeal and eukaryotic homologs that likely represent contamination during genome assembly.**

| Putative archaeal/eukaryotic sequence | Branches with… | Bootstrap support |
|---|---|---|
| *Pantholops hodgsonii*<br>[Tibetan antelope]<br>(XP_005980042.1) | *Asticcacaulis taihuensis*<br>[Proteobacteria]<br>(WP_090646987.1) | 88% |
| *Pantholops hodgsonii*<br>[Tibetan antelope]<br>(XP_005982097.1) | Ochrobactrum sp. PW1<br>[Proteobacteria]<br>(BBA74293.1) | 59% |
| *Aspergillus sclerotialis*<br>(RJE17645.1) | Methylobacterium sp. 4-46<br>[Proteobacteria]<br>(WP_012330955.1) | 49% |
| *Nephila clavipes*<br>[Golden silk orb-weaver]<br>(PRD22299.1) | Acinetobacter<br>[Proteobacteria]<br>(WP_086210128.1) | 96% |
| Thorarchaeota archaeon SMTZ1-83<br>(KXH76530.1) | *Filimonas lacunae*<br>[Bacteroidetes]<br>(SIT08387.1) | 30% |
| Lokiarchaeota archaeon CR 4<br>(OLS13623.1) | *Ktedonobacter racemifer*<br>[Cyanobacteria]<br>(WP_007906963.1) | 97% |

**Table S3. Fourier filtering parameters.**

| **Species** | **Condition** | **Fragment size (bp)** | **PcVal** | **Thr (Zsc)** | **Number of peaks** |
|---|---|---|---|---|---|
| *T. acidophilum* | day1 | 40-65 | 0.02 | 0.25 | 11754 |
| *T. acidophilum* | day2 | 40-65 | 0.02 | 0.25 | 13925 |
| *T. acidophilum* | day3 | 40-65 | 0.02 | 0.25 | 14069 |
| *T. acidophilum* | day3.5 | 40-65 | 0.02 | 0.25 | 8441 |
| *T. acidophilum* | day1 | 70-100 | 0.0125 | 0.25 | 3359 |
| *T. acidophilum* | day2 | 70-100 | 0.0125 | 0.25 | 6887 |
| *T. acidophilum* | day3 | 70-100 | 0.0125 | 0.25 | 6862 |
| *T. acidophilum* | day3.5 | 70-100 | 0.0125 | 0.25 | 4472 |
| *M. fervidus* | Exponential phase | 60-70 | 0.02 | 0.25 | 5363 |
| *M. fervidus* | Exponential phase | 87-97 | 0.0125 | 0.25 | 4780 |
| *M. fervidus* | Exponential phase | 117-127 | 0.01 | 0.25 | 3694 |
| *T. kodakarensis* | Exponential phase | 55-65 | 0.02 | 0.25 | 8472 |

**Supplementary Text**

*A phylogeny of HU proteins and the origin of HTa*

In this supplementary note, we will elaborate on the phyletic distribution of HU proteins outside bacteria and the origin of HTa in the Thermoplasmatales/DHVE2 clade.

First, the hits to archaeal (N=30) and eukaryotic (N=164) genomes that we obtained from structure-guided homology searches (see Materials and Methods), arguably fall into two classes. The first class comprises putative homologs that are isolated from other archaeal/eukaryotic hits on the phylogeny in Figure 2. In principle, these putative homologs might constitute rare cases of horizontal gene transfer (HGT), with a narrow phylogenetic footprint indicating recent arrival. In many, if not most instances, however, these cases likely represent bacterial contaminants in published genome assemblies. This particularly concerns a number of cases where purportedly eukaryotic/archaeal sequences branch with high support with a bacterial sequence. Some examples of such pairs are provided in Table S2. Further indicating likely contamination, several phylogenetically haphazard hits are found even in, for example, mammalian genomes, where rates of horizontal transfer are thought to be extremely low. A striking example is the genome of the Tibetan antelope (*Pantholops hodgsonii*), which – at face value – harbours three different HU proteins affiliated with divergent branches of the bacterial HU phylogeny (Figure 2, Table S2). Contamination might also be the most conservative (and parsimonious) explanation for some hits to phylogenetically isolated archaeal genomes, especially when these were assembled from metagenomic samples (Figure 2, Table S2).

Hits belonging to the second class, in contrast, have at least some phylogenetic persistence and coherence, indicative of vertical inheritance. Amongst eukaryotes, this notably includes homologs in dinoflagellates, some algae, and apicomplexa (including Plasmodium, Theileria, and Babesia species) – all single-celled organism, which have either acquired HU proteins from their resident organelles or unrelated

HGT events. As mentioned in the main text, functional roles for HU proteins have been described for at least some species in these clades.

There are few putative HU homologs in archaeal genomes, with the majority of hits (23/30) belonging to the Thermoplasmatales/DHVE2 clade. Thermoplasmatales/DHVE2 hits are monophyletic (85% bootstrap support, Figure 2), with the exception of a single sequence (Thermoplasmata M8B2D, PNX46291.1, Figure 2), which clusters with 71% bootstrap support with HU from *Desulfobacula toluolica*, a proteobacterium, potentially indicative of contamination or recent HGT. A small number of sequences (4) from halophilic archaea, including Halorubrum aidingense JCM 13560 (EMA66313.1), are embedded amongst Bacteriodetes sequences and appear to form a reasonably coherent phylogenetic unit (Figure S9). Note, however, that the sequence search space contained many other genomes from halophilic archaea, including model organisms such as *Haloferax volcanii,* which did not yield any hits, suggesting that HU proteins in these four species might, again, represent recent acquisitions or assembly contaminants rather than established parts of the functional genome.

Regarding the likely origin of HTa in the Thermoplasmatales/DHVE2 clade, we note that relationships amongst deeper nodes are generally very poorly resolved (as illustrated by smaller node sizes in Figure 2), including in relation to the Thermoplasmatales/DHVE2 clade. There is no strongly supported relationship with particular bacterial sequences that would suggest an ancestral bacterial donor clade. However, HGT remains the most parsimonious explanation for the phyletic distribution of hits, especially in light of the observation that HTa is absent from other members of the larger Diaforarchaea clade, within which Thermoplasmatales/DHVE2 branch.
